# Dietary intervention improves health metrics and life expectancy of the genetically obese DU6 (Titan) mouse

**DOI:** 10.1101/2020.05.11.088625

**Authors:** Annika Müller-Eigner, Adrián Sanz-Moreno, Irene de-Diego, Anuroop Venkateswaran Venkatasubramani, Martina Langhammer, Raffaele Gerlini, Birgit Rathkolb, Antonio Aguilar-Pimentel, Tanja Klein-Rodewald, Julia Calzada-Wack, Lore Becker, Sergio Palma-Vera, Benedikt Gille, Ignasi Forne, Axel Imhof, Chen Meng, Christina Ludwig, Franziska Koch, Angela Kuhla, Vanessa Caton, Julia Brenmoehl, Jennifer Schoen, Helmut Fuchs, Valerie Gailus-Durner, Andreas Hoeflich, Martin Hrabe de Angelis, Shahaf Peleg

## Abstract

Suitable animal models are essential for translational research, especially in the case of complex, multifactorial conditions, such as obesity. The outbred mouse line Titan (DU6) results from the world’s longest selection experiment for high body mass and was previously described as a model for metabolic healthy (benign) obesity. The present study deeper characterized the geno- and phenotypes of this outbred mouse line and tested its suitability as an interventional obesity model. In contrast to previous findings, our data suggests that Titan mice are metabolically unhealthy obese and short-lived. Line-specific patterns of genetic invariability are in accordance with observed phenotypic traits. Titan mice show modifications in the liver transcriptome, proteome and epigenome that are linked to metabolic (dys)regulations. However, dietary intervention partially reversed the metabolic phenotype in Titan mice and significantly extended their life expectancy. Therefore, the Titan mouse line is a valuable resource for translational and interventional obesity research.

## Introduction

Animal models are essential tools for translational biomedical research. The identification of suitable animal models is critical for the successful translatability of pre-clinical data^1–3^. Each animal model presents limitations and that the use of complementing models might highlight particular aspects of a disease. This is critical to model complex, multifactorial diseases, such as metabolic disorders and obesity. The genetic homogeneity of inbred lines, for instance, represents both a strength and a limitation for animal models. While they are useful experimentally by limiting the inter-animal variability, inbred lines might not reflect the genetic heterogeneity characterizing some human diseases^1–3^. A model based on the selection of phenotypic traits (e.g. obesity) may provide novel insight into complex disorders^1,4^.

An example of such selection line is the Dummerstorf “Titan” mouse. Titan (also called DU6) is a unique mouse model resulting from an ongoing long-term breeding scheme (currently over 180 generations) that selected mice based on high body mass at six weeks of age^5^. The initial population of mice was created in 1969/1970 by crossing four outbred and four inbred populations^6,7^ creating a non-inbred mouse line (FZTDU) with a polygenetic background. Based on this polygenetic mouse line, the selection for high body mass started in 1975 with every generation consisted of 60-80 mating pairs^8,9^ (see Methods). Importantly, an unselected population was maintained as a control group throughout the selection process^9^.

Several phenotypic and molecular analyses of the Titan mice were conducted over the course of the selection experiment^5,10–13^. It was reported that 6-week-old Titan mice from earlier selected generations displayed high body mass, as well as elevated levels of fat, insulin, leptin and growth hormone^5,10,13^. An epididymal fat gene expression study identified 77 differentially expressed genes (between Titan and control mice) involved in regulatory and metabolic pathways^11^. A more recent study showed that Titan mice (generation 146) displayed impaired glucose tolerance six weeks of age, while surprisingly, glucose tolerance progressively improved with age compared to unselected control mice^5^. Within the scope of this observation, the Titan (DU6) mice have been proposed as a mouse model to specifically study benign obesity^5^. While metabolically healthy obesity (MHO) is characterized by preserved insulin sensitivity, further characteristics are important to distinguish between MHO to metabolically unhealthy obesity (MUO)^14^. This includes, but is not limited to, lipid profile, liver function, cardiovascular disease, physical activity and healthy life span^14^. Importantly, various obesity-associated comorbidities develop over time and become more prevalent in the later stages of life. Comprehensive analyses of Titan mice throughout their life are lacking^15^.

In this work, we set out to test the hypothesis that the obese Titan mice developed typical hallmarks of pathological detrimental obesity (MUO)^14^. To address this hypothesis, we conducted a deep phenotypic characterization of the mice and investigated factors involved in detrimental obesity in humans or other model species such as plasma levels of cholesterol and triglycerids^16^, Leptin^16–18^, FGF21^19,20^, heart morphology^21^and whitening of adipose tissue^22^. Since body mass and lifespan are two traits known to be inversely correlated to each other^23^, we also assessed whether the lifespan of Titan mice was shortened compared to control mice. In addition, we used recently published genomic sequence data to link several phenotypic features of the mouse line to their specific genetic differentiation caused by the selection procedure^9^.

Furthermore, since the liver is an organ central to lipid and carbohydrate metabolism, we evaluated possible molecular alterations in liver tissue. Specifically, we examined transcriptomic^24^, epigenetic and proteomic alterations that might be associated with obesity and diet. Importantly, dietary intervention is a well-studied and robust intervention to promote healthy lifespan^25–27^ and to improve obesity phenotypes^4,28^ in various animal models. Since not all mouse strains are responsive to caloric restriction^29,30^, we also tested whether the Titan mice might be amenable to therapeutic intervention, using a late and moderate diet intervention as a proof-of-principle.

Our data support the notion that Titan mice exhibit detrimental obesity (MUO), rather than benign obesity (MHO). Titan mice display genetic variability in genomic regions containing genes associated with several regulatory pathways, notably metabolic regulation. The Titan mice exhibit a shorter lifespan, along with molecular changes in liver. Importantly, dietary intervention in Titan mice partly reversed the obesity, transcriptomics phenotypes and lifespan supporting the idea that Titan mice might be suitable as a translational and interventional model of obesity.

## Results

This study was conducted in male Titan (DU6) mice to avoid sex-specific confounding factors. Titan mice aged between 10 and 21 weeks were used for genomic and phenotypic analyses.

### Titan mice display increased size, hallmarks of metabolically unhealthy obesity and reduced lifespan

#### Weight development, gross morphology and abdominal fat distribution

We first conducted a thorough characterization of the phenotype in order to support or refute the notion that the obesity in Titan is benign (MHO vs MUO)^5^.

We started by revisiting the phenotypic characterization of the Titan mice, now at generation 180 of the breeding scheme. Titan mice had an average weight of 90 g at the time of selection (6 weeks of age) vs. 35 g for unselected control mice (Figure 1A). Adult Titan mice further grew to an average of 110-115 g (Figure 1B) and their body and skeleton were substantially larger than those of controls (Figure 1C). At 10–11 weeks of age, Titan mice reached 14.75 cm in body length (Figure 1D). Notably, Titan mice exhibited an increase in both total fat and lean mass (Figure 1E), as well as in fat percentage (Figure 1F). Fat distribution was also altered, with Titan mice accumulating as much as 4% intra-abdominal fat of total body weight compared to 1% in control mice (Figure 1G). Overall, our data confirmed the large and obese phenotype of the mice but also demonstrate an increase in body mass compared to previous data^5,10^.

**Figure 1.**
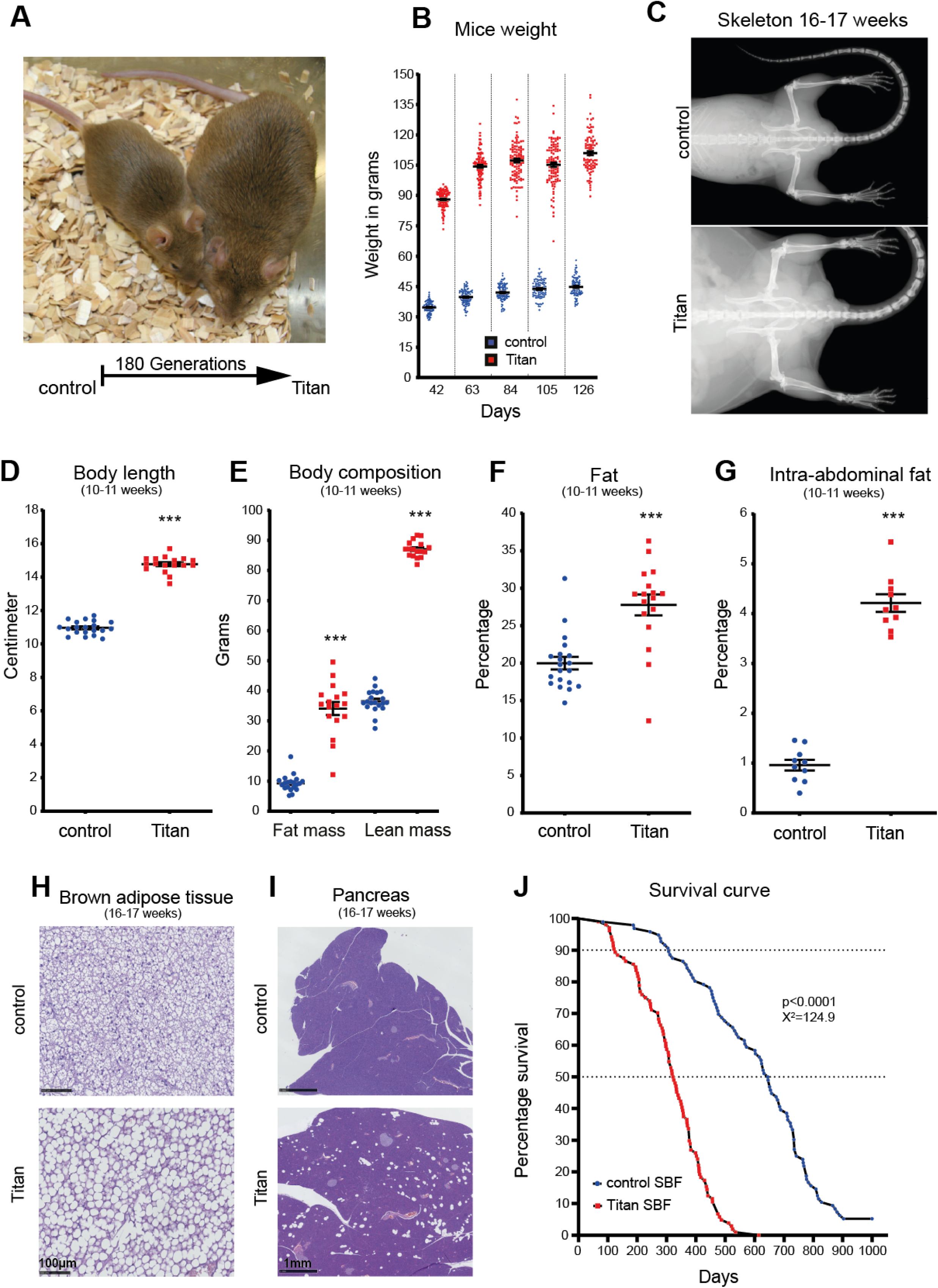
Titan mice, a product of 180 generations of selection, is a severely obese, giant and short-lived mouse strain. (A) Representative images of unselected control mice (left) and 180 generation-selected Titan mice (right) at 6 weeks, at the time of the selection. (B) After the first 16-19 weeks of life, Titan mice reached 110–115 g, whereas average control animals weighed 45 g. (C) Representative X-ray images of control and Titan mice at 16-17 weeks of age. Control (n = 38) and Titan (n = 31) mice. (D) Body length (cm) of 10–11-week-old Titan mice compared to age-matched control mice. Control (n = 20) and Titan (n = 17) mice. (E) Total fat and lean mass of control (n = 20) and Titan (n = 17) mice at the age of 10–11-weeks. (F) Total percentage of fat in control (n = 20) and Titan (n = 17) mice. (G) Percentage of intra-abdominal fat (n = 10 per group) at the age of 10–11-weeks. (H) Representative images of hematoxylin and eosin (H&E)-stained brown adipose tissue (BAT) from control (n = 6) and Titan (n = 5) mice (20x magnification) at 16–17 weeks of age. (I) Representative images of H&E-stained pancreas from control and Titan (n = 6) mice (2.5x magnification) at 16–17 weeks of age. (J) Comparison of lifespans and mortality rates of Titan and control male mice showing significantly reduced lifespan of Titan mice (Log Rank test; p <0.0001, χ^2^=124.9). \*\*\**P* < 0.001. Error bars indicate SEMs. Unpaired two-tailed *t*-tests with Welch’s correction were used to calculate *P*-values.

#### Hallmarks of detrimental obesity

Since obesity is often linked with hyperlipidemia^16^, we measured the cholesterol and triglyceride plasma levels in Titan mice. We detected significantly higher fasting plasma levels of triglycerides, non-HDL and HDL cholesterol in Titan compared to control mice at 18-20 weeks of age (Figure S1A-C). Consistent with the notion that hyperlipidemia usually occurs in the presence of insulin resistance, insulin levels were more than 3-fold higher in ad libitum 16-17-week-old Titan mice (Figure S1D). We also measured average levels of leptin, a hormone secreted by adipocytes and known to modulate food intake and fat storage^16–18^. Leptin levels were significantly higher (more than 5-fold; P < 0.001) in 16-17-week-old Titan mice compared to control mice (Figure S1E). Finally, we found that the plasma concentration of fibroblast growth factor 21 (FGF21), a factor associated with metabolic syndrome (MetS) in humans^19,20^, was elevated in the serum (Figure S1F).

As metabolically unhealthy obesity (MUO) can cause heart disease, we also assessed histological alterations in the heart of the Titan mice^14,21^. Using Sirius Red staining, we detected fibrotic tissue in the hearts of three out of six 16-17-week-old and five out of six 24-26-week-old Titan mice (Figure S2). Furthermore, whitening of adipose tissue has been associated with obesity and obesity-related diseases^22^. Indeed, at 16-17 weeks of age, Titan mice showed a substantial whitening of brown adipose tissue (BAT), not observed in control animals (Figure 1H). In addition, Titan mice showed multifocal fatty cells in the pancreas, termed pancreatic lipomatosis or steatosis, without atrophic or inflammatory changes of the adjacent parenchyma, a phenotype not presented in control mice (Figure 1I).

The lifespan of Titan mice was dramatically shorter than that of control mice (Log Rank test; p<0.0001, χ^2^=124.9) (Figure 1J). Under Standard Breeding Food (SBF), Titan mice reached 10% and 50% mortality at 125 and 325 days, respectively. Their average and maximal lifespans were 317.5 and 614 days (Table S3). In comparison, control mice showed double mean lifespan, reaching 10% and 50% death at 307 and 645 days, respectively. The shortened lifespan of Titan mice combined with obese metrics suggest that the Titan mice suffer from MUO rather than exhibiting MHO.

### Association between phenotypic features of Titan mice and regions of distinct genetic differentiation

Based on the phenotypic analysis, we hypothesized that several genetic alterations in Titan mice would translate to the observed phenotype (metabolic regulation, weight and size). We first assessed the genetic differentiation associated with the Titan mouse, in comparison to the control unselected line but also to other Dummerstorf long-term selected mouse lines^9^. For this, we used the whole genome sequencing data recently generated from the ‘Dummerstorf mouse lines’^9^. We first re-analyzed regions of distinct genetic differentiation (RDD) unique to Titan mice and identified 84 genes (Table S1). Since both Titan and DU6P are giant mice (see ref ^9^), an overlap of RDD was expected between both lines. A second analysis excluding the DU6P mice was conducted and identified 173 protein coding genes (335 including non-coding genes) in RDD for Titan mice (Table S2). Gene Ontology (GO) enrichment analysis (Biological Process) of these Titan-specific RDD genes revealed an association with several biological processes, including metabolic regulation (*Kat2a*, *Sfrp5*, *Agpat4* and *Hcrt*), growth factor signaling (*Stat3*, *Stat5a*, *Stat5b* and *Igf2r*), skin differentiation and immune regulation (Table 1).

**Table 1.**

| *geneSet* | *description* | *size* | *overlap* | *expect* | *enrichmentRatio* | *pValue* | *FDR* | *overlapId* | *userId* |
| --- | --- | --- | --- | --- | --- | --- | --- | --- | --- |
| GO:0031424 | keratinization | 35 | 5 | 0.2304330 | 21.698286 | 0.0000032 | 0.0196906 | 20754; 20762; 20763; 20765; 20766 | Sprr2h; Sprr2i; Sprr2k; Sprr1b; Sprr3 |
| GO:0044262 | cellular carbohydrate metabolic process | 285 | 10 | 1.8763826 | 5.329403 | 0.0000193 | 0.0504800 | 11677; 12015; 14042; 14534; 15531; 20848; 50798; 50873; 58246; 66646 | Rpe; Gne; Slc35b4; Akr1b3; Kat2a; Stat3; Ext1; Prkn; Ndst1; Bad |
| GO:0060343 | trabecula formation | 27 | 4 | 0.1777626 | 22.501926 | 0.0000279 | 0.0504800 | 14225; 15214; 21687; 170826 | Fkbp1a; Tek; Hey2; Ppargc1b |
| GO:0060416 | response to growth hormone | 28 | 4 | 0.1843464 | 21.698286 | 0.0000324 | 0.0504800 | 20848; 20850; 20851; 227231 | Cps1; Stat5b; Stat5a; Stat3 |
| GO:0009913 | epidermal cell differentiation | 199 | 8 | 1.3101759 | 6.106050 | 0.0000522 | 0.0650244 | 15214; 20754; 20762; 20763; 20765; 20766; 22283; 67694 | Ush2a; Sprr2h; Sprr2i; Sprr2k; Sprr1b; Sprr3; Ift74; Hey2 |
| GO:0010918 | positive regulation of mitochondrial membrane potential | 13 | 3 | 0.0855894 | 35.051077 | 0.0000759 | 0.0675235 | 12015; 50873; 269523 | Vcp; Prkn; Bad |
| GO:0072044 | collecting duct development | 13 | 3 | 0.0855894 | 35.051077 | 0.0000759 | 0.0675235 | 11677; 71228; 240025 | Akr1b3; Dlg5; Dact2 |
| GO:0060396 | growth hormone receptor signaling pathway | 14 | 3 | 0.0921732 | 32.547429 | 0.0000962 | 0.0744756 | 20848; 20850; 20851 | Stat5b; Stat5a; Stat3 |
| GO:0060347 | heart trabecula formation | 15 | 3 | 0.0987570 | 30.377600 | 0.0001196 | 0.0744756 | 14225; 15214; 21687 | Fkbp1a; Tek; Hey2 |
| GO:0045579 | positive regulation of B cell differentiation | 15 | 3 | 0.0987570 | 30.377600 | 0.0001196 | 0.0744756 | 12015; 20850; 20851 | Stat5b; Stat5a; Bad |
| GO:0046629 | gamma-delta T cell activation | 16 | 3 | 0.1053408 | 28.479000 | 0.0001465 | 0.0751777 | 20850; 20851; 270152 | Jaml; Stat5b; Stat5a |
| GO:0071378 | cellular response to growth hormone stimulus | 16 | 3 | 0.1053408 | 28.479000 | 0.0001465 | 0.0751777 | 20848; 20850; 20851 | Stat5b; Stat5a; Stat3 |
| GO:0050891 | multicellular organismal water homeostasis | 42 | 4 | 0.2765195 | 14.465524 | 0.0001651 | 0.0751777 | 11677; 20276; 22393; 229574 | Flg2; Wfs1; Akr1b3; Scnn1a |
| GO:0035966 | response to topologically incorrect protein | 124 | 6 | 0.8163910 | 7.349419 | 0.0001711 | 0.0751777 | 22393; 50873; 56323; 69310; 69601; 269523 | Dab2ip; Dnajb5; Vcp; Wfs1; Pacrg; Prkn |
| GO:0072528 | pyrimidine-containing compound biosynthetic process | 43 | 4 | 0.2831033 | 14.129116 | 0.0001812 | 0.0751777 | 29807; 68262; 69692; 227231 | Cps1; Tpk1; Hddc2; Agpat4 |
| GO:0030216 | keratinocyte differentiation | 131 | 6 | 0.8624776 | 6.956702 | 0.0002306 | 0.0863868 | 20754; 20762; 20763; 20765; 20766; 67694 | Sprr2h; Sprr2i; Sprr2k; Sprr1b; Sprr3; Ift74 |
| GO:0030104 | water homeostasis | 46 | 4 | 0.3028547 | 13.207652 | 0.0002359 | 0.0863868 | 11677; 20276; 22393; 229574 | Flg2; Wfs1; Akr1b3; Scnn1a |
| GO:0045838 | positive regulation of membrane potential | 19 | 3 | 0.1250922 | 23.982316 | 0.0002499 | 0.0864356 | 12015; 50873; 269523 | Vcp; Prkn; Bad |
| GO:0030856 | regulation of epithelial cell differentiation | 136 | 6 | 0.8953966 | 6.700941 | 0.0002824 | 0.0925190 | 12015; 15214; 20850; 20851; 21937; 74123 | Tnfrsf1a; Hey2; Stat5b; Stat5a; Foxp4; Bad |
| GO:0061383 | trabecula morphogenesis | 49 | 4 | 0.3226061 | 12.399020 | 0.0003017 | 0.0938981 | 14225; 15214; 21687; 170826 | Fkbp1a; Tek; Hey2; Ppargc1b |
| GO:0043588 | skin development | 259 | 8 | 1.7052038 | 4.691521 | 0.0003204 | 0.0949844 | 20754; 20762; 20763; 20765; 20766; 67694; 229574; 240025 | Sprr2h; Sprr2i; Sprr2k; Sprr1b; Sprr3; Flg2; Ift74; Dact2 |

Next, we sought to further explore the link between genotype and phenotype. Increased skin thickness has been shown to positively correlate with increased BMI^31^. As expected, Titan mice displayed a thicker dermis of the skin compared to the control group (Figure S3A). We also found that the inflammatory cytokines interleukin 6 (IL-6) and tumor necrosis factor-alpha (TNFα) were significantly increased in the plasma of Titan mice compared to unselected control mice (Figure S3B and C). As a further indication of inflammation, histological analysis of the thymus revealed the presence of nodular thymic medullary hyperplasia in Titan mice (Figure S3D). Immunohistochemistry of these medullar nodes revealed that they were composed mainly of B cells (Figure S3D). Thus, our data supports a link between Titan-specific RDD such as metabolic regulation, growth, skin differentiation and immune regulation genes with Titan-specific phenotypes of obesity, size, thicker dermis and increased hallmarks of Inflammation.

### Altered histone 4 acetylation levels in Titan mice

Genetics and epigenetics functions are intimately linked, and their alterations are associated with disease^32^. We identified the histone acetyl transferase *kat2a* (*Gcn5*), as one of the genes associated with specific genetic differentiation in Titan mice (Figure 2A, Table S1; Allele frequency = 0.97). Kat2a is a histone acetyl-transferase, which has also been implicated in metabolic and energy regulation^33^, thus raising the possibility that histone acetylation might be altered in Titan mice. To address this possibility, we compared histone 4 (H4) acetylation levels of 10-week-old and 19-20-week-old Titan vs. control mice by mass spectrometry^34,35^. For this analysis, we focused on the liver as it is an organ central to lipid and carbohydrate metabolism and KAT2a is involved in regulation of metabolic activity in the liver^36^.

**Figure 2.**
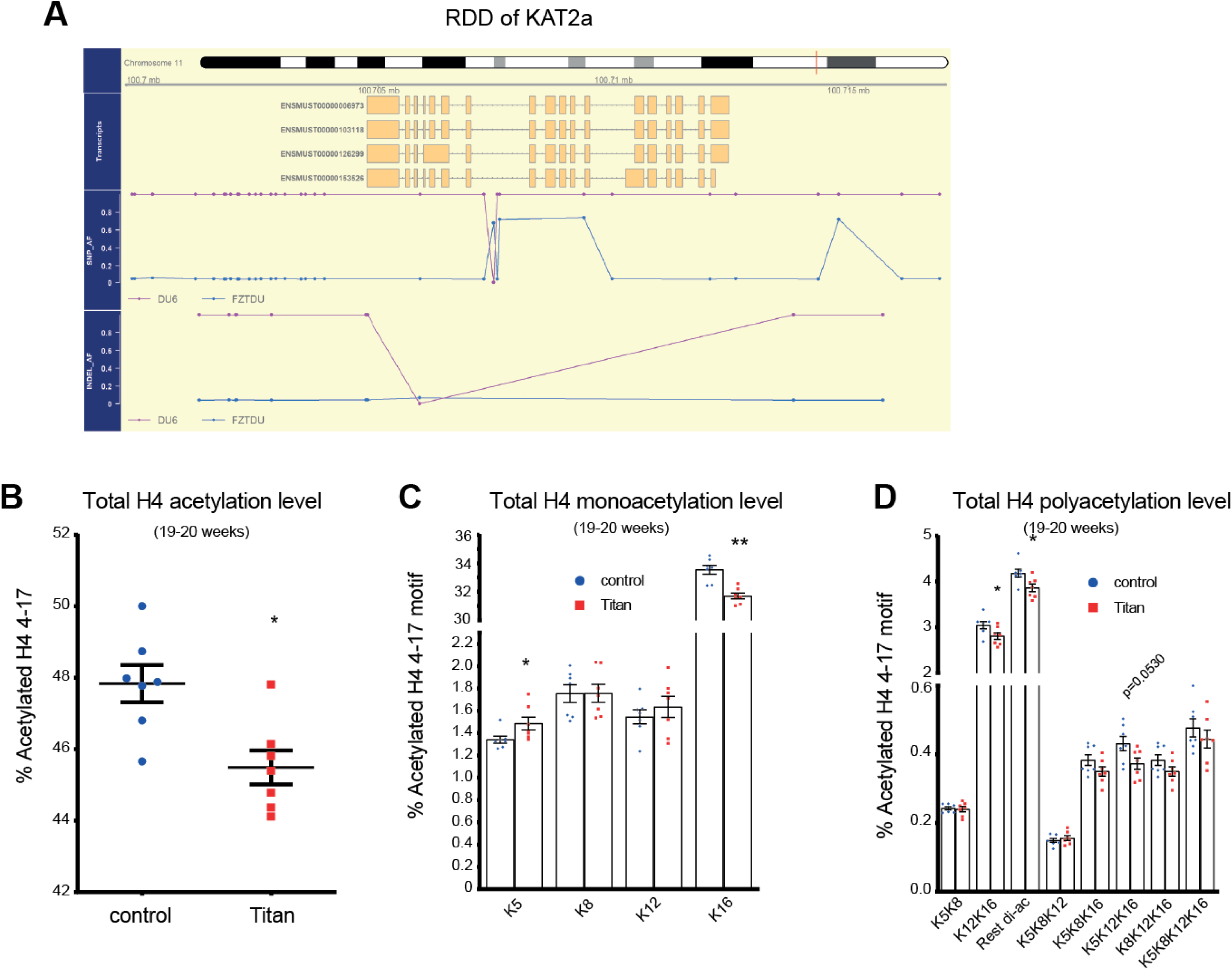
Histone 4 acetylation levels are altered in Titan compared with control unselected mice. (A) Scheme of the RDD of the *kat2a* gene in Titan and control mice. (B) Total levels of histone 4 4-17 levels via mass spectrometry (p =0.0111, MWU-test) (n = 7 per group). (C) Quantification of monoacetylation of histone 4 4-17. P-values, MWU-test (H4K5K8)=0.0175, (H4K8)=0.901, (H4K12)=0.455, (H4K16)=0.0023 (n = 7 per group). (D) Quantification of polyacetylation of histone 4 4-17. P-values, MWU-test (H4K5K8)=1, (H4K12K16)=0.0379, (H4-rest di-ac)=0.0379, (H4K5K8K12)=0.62, (H4K5K8K16)=0.208, (H4K5K12K16)=0.053, (H4K8K12K16)=0.208, (H4 tetra-ac)=0.62 (n = 7 per group). \**P* < 0.05, \*\**P* < 0.01. Error bars indicate SEMs.

At 19-20 weeks of age, we observed significantly lower levels of H4 acetylation in Titan vs. control mice (Fig 2B). Highly abundant H4 acetylation motifs in mammalian such as H4K16 and H4K12K16^35^, as well as poly-acetylated motifs were significantly reduced in Titan vs. control mice (Fig 2C,D). By contrast, H4K5 acetylation motif, which was shown to be moderately regulated at least in part by Kat2a^34^, was significantly higher in Titan mice compared to control mice (Fig 2C). Of note, comparable results were obtained in younger Titan mice (10 weeks of age, data not shown). Taken together, Titan mice display considerable alterations in various histone acetylation motifs compared to the unselected mouse line which might manifest in altered transcriptomes^37^.

### Hybrids of Titan and unselected mice show intermediate phenotypes

We next assessed the potential contribution of the maternal impact and phenotypic features of Titan mice. In order to address this we generated hybrids by crossing Titan and control unselected mice (mother control/Titan x father Titan/control respectively). Male F1 hybrid of 10-11 weeks of age showed similar weight and fat % independently of the genotype of the mother (Fig 3A).

**Figure 3.**
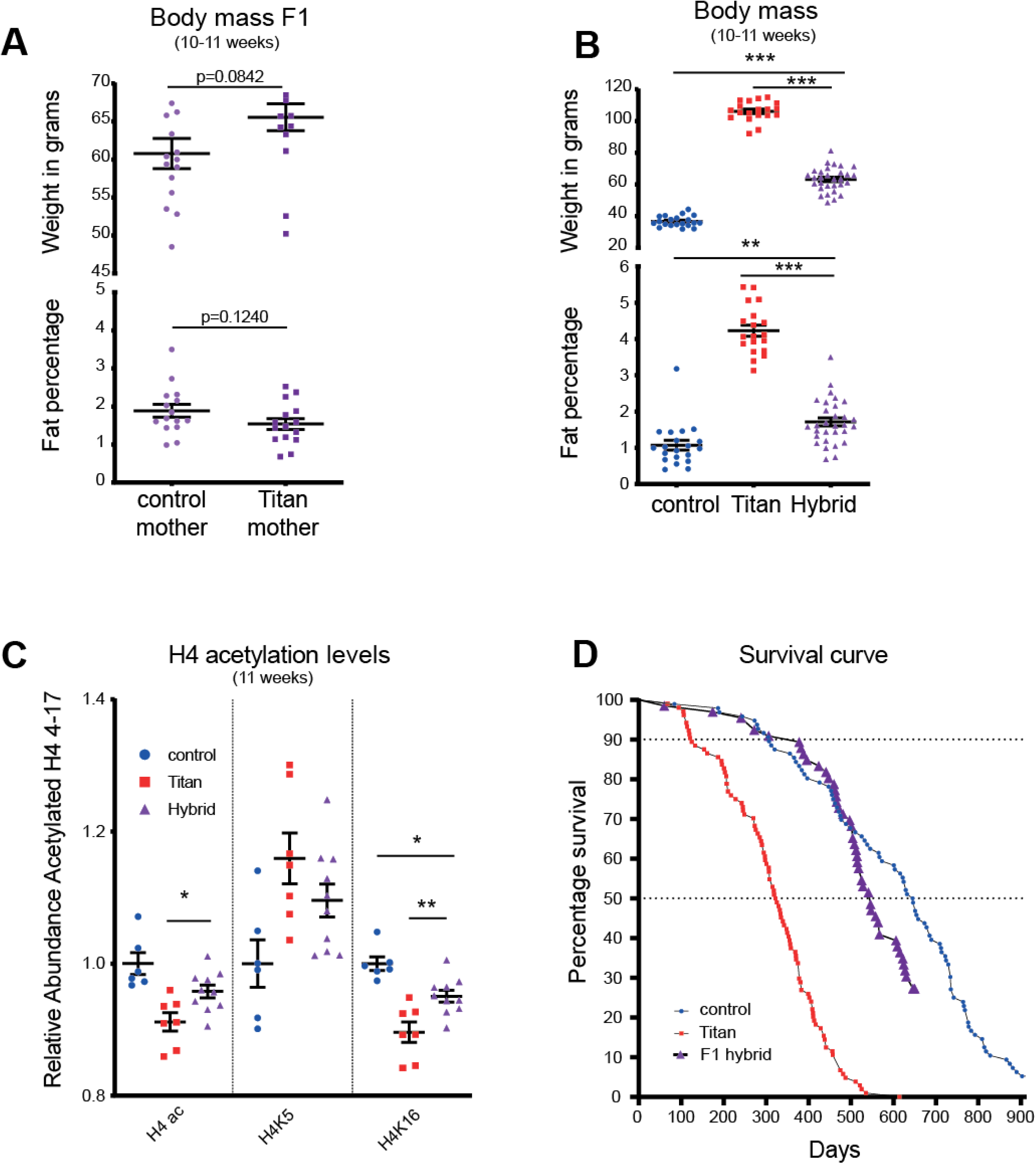
Hybrid offsprings of control and titan mice display intermediate obesity and life span. (A) Body weight and visceral fat percentage in F1 hybrid (Offsprings of either mother control x father mother vs mother titan x father control) n = 15 each, MWU-test was used to calculate p-value.(B) Quantification of total histone 4 4-17 acetylation, H4K5ac and H4K16ac in control (n = 6), Titan (n = 7) and hybrid (n = 10) mice. One way ANOVA followed by Tukey test P-value: control vs hybrid (H4ac)=0.0765 (H4K5)=0.116, (H4K16)=0.0184 and Titan vs hybrid (H4ac)=0.0383, (H4K5)=0.330, (H4K16)=0.0065. (C) Body weight and visceral fat percentage in control (n = 20), Titan (n = 19) and hybrid (n = 30). One way ANOVA followed by Tukey test P-value: control vs hybrid (Weight)<0.0001, (%visceral fat)=0.0021 and Titan vs hybrid (Weight)<0.0001 (%visceral fat)<0.0001. (D) Comparison of lifespans and mortality rates of Titan and control male mice to Hybrid (N=66). \**P* < 0.05, \*\**P* < 0.01. \*\*\**P* < 0.001. Error bars indicate SEMs.

Notably, the combined population of male F1 hybrids showed intermediate body weight and fat percentage compared to control and Titan mice at the same age (Figure 3B). Similarly, the hybrid mice showed intermediate H4 acetylation levels in the liver, including at H4K5 and H4K16 (Figure 3C). Furthermore, the lifespan of hybrid mice, although improved compared to Titan mice, was reduced compared to that of control mice (Hybrid vs Titan Log Rank test; p <0.0001, χ^2^=96.45 Control vs hybrid, Log Rank test; p =0.0362, χ^2^=4.385). The median life span of 546 days vs 325 (Titan) and vs 645.5 days (control)) (Figure 3D). These observations illustrate the quantitative genetic background determining mice obesity and life span.

### Titan mice display altered liver metabolism along with changes in gene and protein expression

We next evaluated possible alterations in liver function and metabolism in Titan mice. First, alanine aminotransferase (ALAT) and alkaline phosphatase (ALP) activity, markers of potential liver damage^38^, were significantly elevated in Titan mice at 16-17 weeks of age (Figure 4A and B). Combined Oil red O and Hematoxylin staining of the liver also revealed numerous small fat droplets in Titan mice (Figure 4C). However, no clear signs of hepatic steatosis were found by H&E staining of 16-17-week-old Titan liver tissue (Figure S4). Bilirubin, a waste product of heme metabolism in the liver, was significantly lower in Titan mice compared to controls (Figure 4D). In addition, plasma urea levels were lower in Titan than in control mice (Figure 4E), pointing to the dysregulation of the urea cycle and protein metabolism in the liver. Altogether, these findings suggest an altered liver metabolism in Titan compared to control unselected mice.

**Figure 4.**
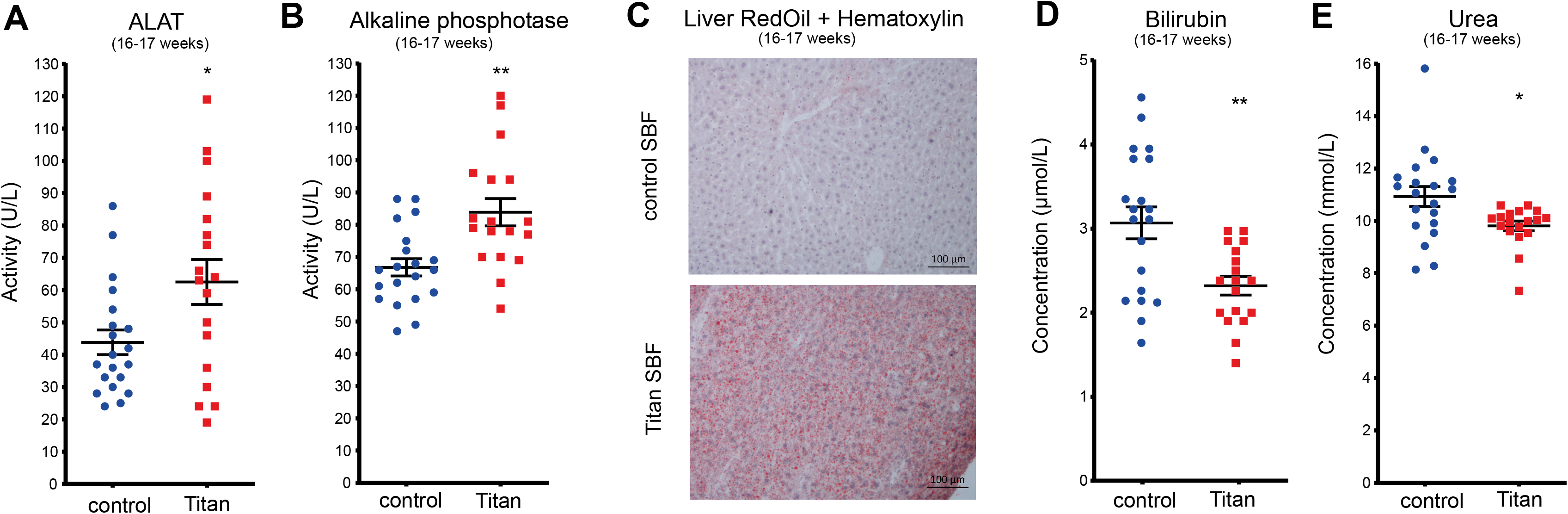
Titan mice display altered liver metabolism. (A) Alanine aminotransferase (ALAT) (p = 0.0262) and (B) alkaline phosphatase (p = 0.0019) levels in control (n = 20) and Titan mice (n = 18). (C) Oil Red O with Hematoxylin staining of fat in liver of control and Titan mice (n = 4 per group). (D) Bilirubin levels in control (n = 20) and Titan mice (n = 18) (p = 0.0019). (E) Urea levels of control (n = 20) and Titan (n = 18) mice (p = 0.0132). \**P* < 0.05, \*\**P* < 0.01. Error bars indicate SEMs. Unpaired two-tailed *t-*tests with Welch’s correction were used to calculate *P-*values.

We hypothesized that Titan mice, unlike control mice, would display transcriptome profiling that is linked to increased body fat phenotype and altered metabolism. To identify pathways altered at the transcriptional level in the liver, we performed RNA sequencing on the liver from 11-week-old and 19-21-week-old mice. Principal component analysis (PCA) revealed distinct Titan and control transcriptomes (Figure 5A). Gene expression in younger and older controls clustered together, but in Titan mice, the expression profile shifted between the age groups (Figure 5A), indicating age-dependent effects in Titan but not in control mice.

**Figure 5.**
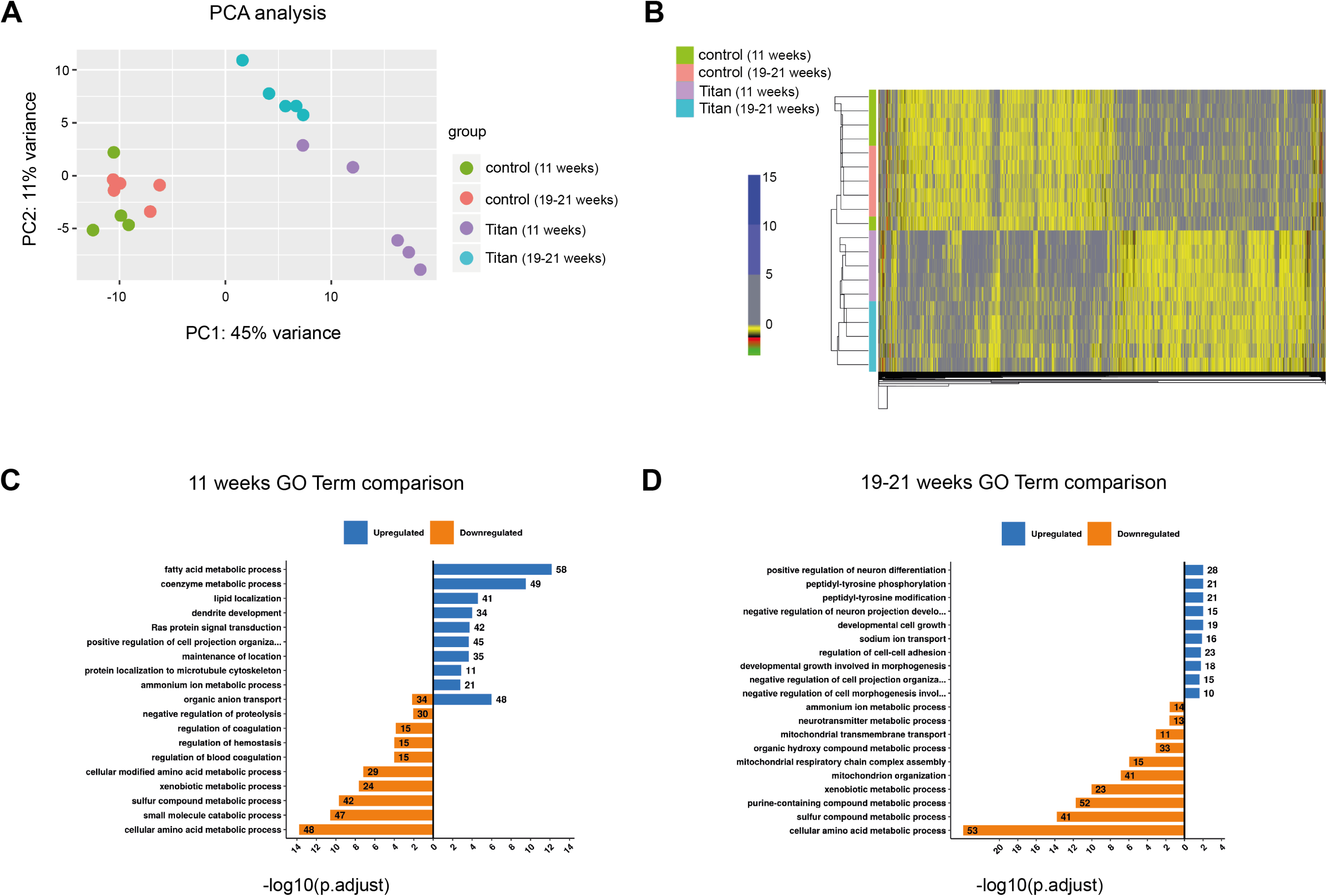
Liver transcriptome analyses reveal differential expression of genes associated with metabolic pathways in Titan mice. (A) Principal component analysis (PCA) of liver transcriptomes revealed clustering differences between Titan mice and controls. Control mice at 11 weeks and 19-21 weeks had similar transcriptomes, while younger and older Titan mice had distinct transcriptomes. (B) Heat map showing significantly altered genes in control and Titan mice at 11 weeks and 19-21 weeks of age. Color gradient in each cell in the heatmap represents the scaled normalized log counts from each replicate and condition. (C) GO term analysis in 11-week-old Titan mice compared with corresponding control. (D) GO term in 19-21-week-old Titan mice compared with corresponding control. (n = 5 per group).

Compared to age-matched controls, 11-week-old Titan mice exhibited 1150 upregulated and 1227 downregulated genes (Figure 5B, S5A and Table S4) and 19-21-week-old Titan mice 693 upregulated and 828 downregulated genes (Figure 5B and S5D). We observed a significant overlap of transcriptomic alterations between both age group comparisons, with 402 upregulated and 461 downregulated genes.

We next compared 11-week-old Titan to control mice. The top 10 upregulated GO terms highlighted genes involved in several central metabolic processes (Figure 5C). Central metabolic regulators such as *mTOR, Elovl5, Elovl6*, and acetyl-CoA carboxylase were upregulated (Table S4), suggesting altered fat metabolism in the liver. The top 10 downregulated GO terms included xenobiotic, amino acid and sulfur metabolism (Figure 5C). For example, many cytochrome P450 family genes, including *Cyp2c37,* were downregulated in 11-week-old Titan mice compared to controls at the same age (Table S4). Interestingly, cytochrome P450 family genes are downregulated in a high-fat diet^24^. Furthermore, several genes related to the methionine, folate cycle and H_2_S production [e.g., *Bhmt, Gnmt, Cth* (*Cgl*)*, Dmgdh*], as well as *Hcrtr2* were downregulated in 11-week-old Titan mice.

The top 10 upregulated GO terms in 19-21 week-old Titan mice compared with control revealed the upregulation of genes involved in neuronal differentiation and peptidyl-tyrosine phosphorylation activity. (Figure 5D). The top 10 downregulated GO terms in 19-21-week-old Titan mice revealed genes involved in the regulation of xenobiotic, amino acid and sulfur metabolism (Figure 5D), similar to the comparison at 11 weeks of age (Figure 5C). Of note, genes involved in mitochondrial organization and mitochondrial respiratory complex assembly were downregulated as well (Figure 5D). We found that *Acss2*, an acetyl-CoA synthesis enzyme linked with histone acetylation^39^, was downregulated in Titan mice.

Compared to 11-week-old Titan mice, 19-21-week-old Titan mice had increased expression of several enzymes involved in glucose and insulin response (See supplementary data). Also, *Cyp7b1*, which encodes an enzyme involved in cholesterol catabolism that converts cholesterol to bile acids and is upregulated by high-fat diet^24^, was upregulated in 19-21-week-old Titan mice compared to 11-week-old Titan mice. On the other hand, the top 10 downregulated GO term highlighted genes involved in fatty acid, coenzyme and carbohydrate metabolism (Figure S5C). Taken together, liver transcriptomic analysis demonstrate several metabolic alterations in Titan mice.

Next, we compared the liver proteomes of Titan and control mice. As with the RNAseq data, larger proteomic changes were seen in the 11-week-old Titan to control comparison than in the 19-21-week-old comparison (Figure S6). Relative to age-matched controls, 11-weeks-old Titan mice exhibited 207 upregulated and 289 downregulated proteins, while 19-21-week-old Titan mice showed 125 upregulated and 142 downregulated proteins (Figure S6). Consistent with the transcriptome results, pathway analysis revealed increased fatty acid metabolism, lipid catabolism, and coenzyme metabolism, notably pathways involved in fat utilization (ACACA, IDH1 and ACOX1) and in lipid biosynthesis (FASN and ACSL5), while downregulation in xenobiotic, amino acid and sulfur metabolism in 11-week-old Titan mice (Figure S6).

Proteins involved in lipid metabolism were also enriched in 19-21-week-old Titan mice compared to controls (Figure S6), although this was not observed in the RNAseq top 10 GO terms (Figure 5D). We also found that proteins upregulated in 19-21-week-old Titan mice were associated with carbohydrate catabolism and coenzyme metabolism (Figure S6), while downregulated proteins were enriched in amino acid and sulfur metabolism (Figure S6), in line with the RNAseq analysis (Figure 5D). As with the transcriptomic data, we observed a reduction in the abundance of BHMT, GNMT, and CTH (CGL).

Collectively, both transcriptomics and proteomics data hint towards increased fat metabolism in Titan mice compared to control mice.

### Late dietary intervention at 12 weeks of age improves the obesity and lifespan phenotypes in Titan mice

Finally, we investigated the impact of dietary restriction on measures of obesity, epigenetic markers, liver gene expression patterns and lifespan of Titan mice. We compared the impact of a diet with moderate energy reduction using Energy Reduced Food, hereafter referred to as ERF (see Methods), to that of Standard Breeding Food (SBF). We introduced ERF at 12 weeks of age onward and termed this diet intervention as late intervention due to the fact that 10% and 25% of the Titan population are already dead by 18 and 33 weeks, respectively.

ERF intervention resulted in a weight reduction of 5–10% in Titan mice, compared to SBF-fed siblings (Figure 6A; MWU-test, P=0.0017). By 21 weeks, ERF-fed Titan mice had significantly reduced abdominal fat percentage compared to SBF-fed mice (Figure 6B; MWU-test, P=0.0313). We also showed that the ERF regime lowered the levels of plasma cholesterol, HDL and glucose in Titan mice (Figure 6C). Furthermore, plasma levels of leptin, glycerol and non-esterified fatty acids (NEFA) were decreased under ERF regime (Figure 6C).

**Figure 6.**
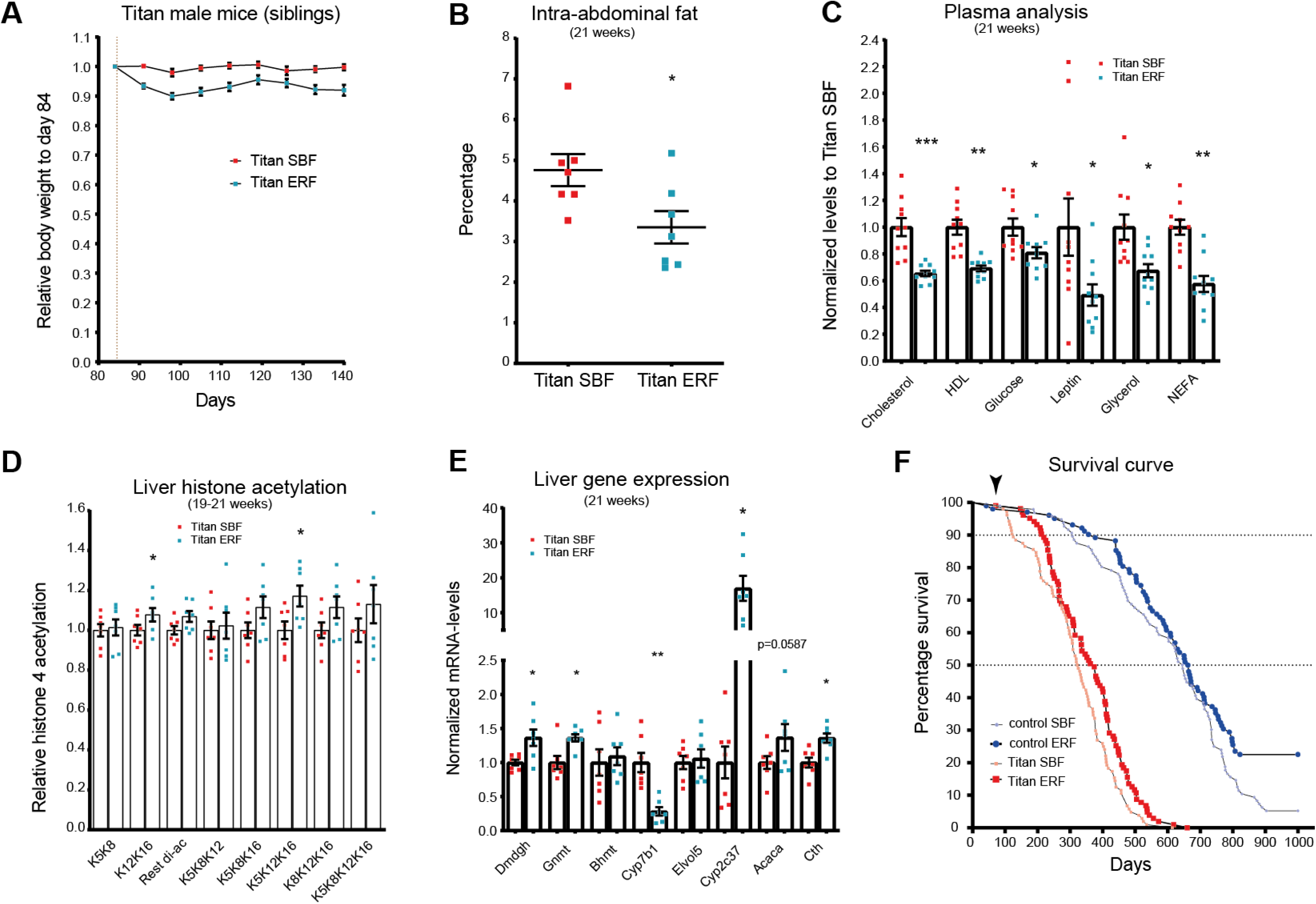
Late dietary intervention by switching to energy-reduced food (ERF) improves health metrics and extends the life span of Titan mice. (A) Switching standard feeding (SBF) to ERF at 12 weeks resulted in a persistent average weight loss in Titan siblings (n = 23 per group). MWU-test, P=0.0017. (B) Compared to age-matched control Titan mice, ERF-fed Titan mice had a lower percentage of intra-abdominal fat at 21 weeks (n= 7 per group). Paired MWU-test, p = 0.0313. (C) Late ERF feeding at 12 weeks of age decreased the levels of plasma cholesterol (pval.adj=0.00013), HDL (pval.adj=0.0013), Glucose (pval.adj=0.043), Leptin (pval.adj=0.043), Glycerol (pval.adj=0.0165) and NEFA (pval.adj=0.00132) in 21 weeks old mice. (n= 10 per group). MWU-test, by multiple correction to calculate P-values. (D) Quantification of polyacetylation of histone 4 4-17 following ERF feeding. P-values, MWU-test (H4K5K8)=0.8125, (H4K12K16)=0.0469 (H4-rest di-ac)=0.1563, (H4K5K8K12)=0.9375, (H4K5K8K16)=0.0781, (H4K5K12K16)=0.0313, (H4K8K12K16)=0.0781, (H4 tetra-ac)=0.375 (n = 7 per group) Paired MWU-test, to calculate P-values. (E) RT-qPCR comparing gene expression of candidate genes (from table S4) of ERF- and control-fed Titan mice siblings at 21 weeks of age (n = 7 per group). Paired t-test was performed followed by multiple correction to calculate P-values pval.adj (*Dmdg*)=0.032, (*Gnmt*)=0.012, (*Bhmt*)=0.736, (*Cyp7b1*)=0.008, (*Elovl5*)=0.736, (*Cyp2c37*)=0.012, (*Acaca*)=0.0587, (*Cth*)=0.012. (F) Switching to ERF feeding at 12 weeks of age (vertical dashed line) increased the lifespan of both the Titan and control mice. The ages at 25%, 75% and 90% death, and the median lifespan are presented in Table S1. *P < 0.05, **P < 0.01, ***P < 0.01. Error bars indicate SEMs.

Late ERF intervention in Titan also resulted in a general increase in poly-acetylated motifs of histone 4 levels, which resembled the levels observed in control mice (Fig. 6D). In general, motifs containing K12K16 acetylation motifs (Significant H4K12K16 and H4K5K12K16) were increased in ERF-fed compared to SBF-fed Titan mice (Figure 6D). We next tested whether the ERF regime affected the expression level of liver genes that were already identified in our omics experiments as significantly altered in Titan compared to control mice (Figures 4 and S6, See supplement list of genes). In addition, late ERF intervention reversed the expression profiles of some of the investigated genes. In fact, genes that were downregulated in Titan versus control mice (e.g. *Dmdgh, Gnmt, Cth, Cyp2c37*) were upregulated in ERF-fed vs SBF-fed Titan mice (Figure 6E)^24^. Similarly, genes which were upregulated in Titan versus control mice (e.g. *Cyp7b1*) were downregulated in ERF-fed vs SBF-fed Titan mice (Figure 6E).

Finally, we investigated the impact of the ERF diet on the lifespan of Titan and control mice. The ERF diet significantly increased the lifespan of both Titan and control mice (Log Rank test; for P=0.0087, χ^2^=6.892 Titan mice and P=0.01, χ^2^=6.66 for control mice) (Figure 6F). ERF-fed Titan mice reached the 10% and 50% death at 219 and 374 days, respectively, which is significantly longer than for the SBF-fed Titan mice (10% death at 125 days and 50% death at 325 days). In comparison, ERF-fed control mice reached 10% death at 377 days vs 307 days for SBF-fed control mice. The average lifespan of ERF-fed Titan mice was 359.5 days, compared to 317.4 days for SBF-fed Titan mice (Table S3).

Taken together, our results indicate that a late ERF intervention could partly revert the phenotypes of Titan mice associated with MUO and lifespan.

## Discussion

This study provides a comprehensive phenotypic analysis of the Titan mouse line DU6 and investigated the link between Titan’s unique phenotype to their genotype. We demonstrate that Titan mice display various physiological and molecular alterations that may underlie their metabolically unhealthy obesity phenotype. We also show that Titan mice are short-lived, and that the obesity and lifespan phenotypes are responsive to and can be partly reversed by diet intervention.

We provide several links between Titan mice’s genotype and their unique phenotypes. Gene Ontology (GO) enrichment of regions of distinct genetic differentiation (RDD) highlighted genetic variability associated with genes involved in multiple pathways including metabolic regulation and growth control. Similarly, RDD genes involved in skin differentiation correlated with our observation that Titan mice display a thicker dermis of the skin compared to control mice, which is in line with previous observations linking increased BMI with thicker skin^31^. Several RDD genes linked to immune regulation and inflammation (notably Stat3 and Stat5 family members) were also identified^40^, in correlation with the immune and inflammatory phenotype of Titan mice. For example, Titan mice expressed markers of systemic inflammation, such as high plasma levels of IL-6 and TNFα or thymic medullary hyperplasia. Besides being associated with the initiation and progression of multiple diseases^41,42^, elevated circulating IL-6 and TNFα were documented in obese mice and humans^43,44^. Supporting this finding, several mouse models of obesity revealed increased accumulation of macrophages in the liver^45,46^ or in adipose tissue^47^. Increased tissue inflammation in Titan mice might contribute to tissue damage^38^.

Notably, one identified RDD contained the *Hcrt* gene, which is a hypothalamic neuropeptide that can regulate feeding behavior and metabolic homeostasis^48,49^. While we did not evaluate gene expression levels in the brain area that may account for metabolic alterations in the Titan mice, we observed a significant reduction of *Hcrtr2* in the liver. Intriguingly, *Kat2a*, which encodes the histone acetyltransferase GCN5, was also part of a RDD specific to Titan mice and is involved in metabolic regulation^33^. This correlated with an increase in H4K5 acetylation in the liver of Titan mice, suggesting a possible causal link between the two phenotypes^34^. However, we did not detect significant changes in the expression level of *Kat2a* in the liver of Titan mice. Although it is unlikely that changes in H4K5 acetylation are the consequence of changes in *Kat2a* expression, we cannot exclude that they are related to changes in *Kat2a* enzymatic activity or to alterations in other GCN5-related genes. Additional investigations will be necessary to address this question and further assess alterations in expression and activity of histone acetyltransferases in Titan mice, in correlation with changes in histone acetylation levels.

Notably, F1 hybrids show intermediate body weight and fat percentage, H4 acetylation levels and lifespan. The intermediate phenotype is independent of the mother’s genotype (Control or Titan), thus demonstrating a weaker link between maternal inheritance/ care to weight and fat levels. Further analyses of F2 hybrids are required to link specific genes to the specific phenotypes of the Titan mice described in this study^4,50^.

Previous research at earlier generations suggested that Titan mice display benign obesity (MHO) without additional comorbidities^5^. In contrast, we now show that Titan mice indeed exhibit detrimental obesity, including several hallmarks that are attributed to metabolic syndrome (MetS), such as high levels of fat in tissues, high plasma levels of fasting triglycerides, fasting cholesterol, FGF21^19,20,51^ and insulin^16,52^ as well as early signs of heart fibrosis^21^. Titan mice also showed whitening of BAT, and ectopic fat in the pancreas, both being linked to obesity and MetS^22,53^. Increased pancreatic fat might have implications for the onset of type 2 diabetes^54^, which is frequently associated with insulin resistance^21^. Importantly, obesity is known to be associated with reduced life expectancy and various conditions related to aging, such as inflammation^55–60^. Indeed, we found that Titan mice exhibited a shorter lifespan compared to control mice, which is in line with a recent study showing that female Titan mice show a reduced lifetime fecundity compared to the unselected control mice^61^.

Alterations in liver function were evidenced by epigenomics, transcriptomic and proteomic changes detected in the liver of Titan mice. Genes and proteins involved in lipid metabolism and biosynthesis were consistently upregulated in Titan mice compared to control mice, which might account for the high plasma levels of cholesterol and fasting triglycerides observed in this model. On the other hand, compared to control mice, Titan mice exhibited several downregulated genes and proteins associated with xenobiotic metabolism, such as cytochrome P450. These downregulations might indicate altered endogenous metabolite metabolism^62^, increased liver inflammation^63,64^ and/or a generally reduced capacity to deal with toxic compounds in the liver. Xenobiotic metabolism strongly impacts the ability of the organism to maintain homeostasis and cope with disease^65^, which may contribute to the increased morbidity of these mice. Also, Titan mice showed a reduced expression of genes related to the folate cycle and its downstream effectors, such as *Ahcy*, *Gsta* family *Gnmt* and *Cth*. Such genes were also shown to be downregulated in the mouse liver upon high-fat diet^24^. Similarly, genes involved in mitochondrial organization and function were downregulated in 19-21-week-old Titan mice, which may hint towards altered mitochondrial activity in older Titan mice^58^. Altogether, these findings suggest an altered liver metabolism in Titan compared to control mice, along with their obese phenotype and shorter lifespan.

Interestingly, the liver of Titan mice showed overall decreased H4 acetylation at the 4-17 motif, specifically histone acetylation motifs such as H4K16 and H4K12K16 acetylation, which is in correlation with a reduction in the expression of acetyl-CoA synthases (*Accs2* and *pdhx*) which may imply reduced acetyl-CoA availability. This may underlie the general alteration in histone acetylation levels^39,66^ and might also link this epigenetics phenotype to lifespan regulation^67^.

Our data support the notion that Titan mice are a potential translational model, as ERF feeding introduced at 12 weeks of age could partially reverse the phenotypes associated with obesity, lifespan and histone H4 acetylation patterns. Notably, intra-abdominal fat was reduced at 21 weeks and plasma levels of leptin, glycerol and non-esterified fatty acids (NEFA), which are linked to obesity^16,17,68,69^, were decreased in ERF-fed Titan mice. This phenotype improvement is reminiscent of other time-restricted fasting animal models or patients with metabolic syndrome^70^. ERF feeding appears to specifically increase levels of poly acetylated histone 4 motif which include K12K16 acetylation. These modification might impact transcription regulation that are important for mediating the benefits of healthy diet in obese animals and maybe novel therapeutic targets to treat obesity. In line with these data, we also found that ERF feeding partially reversed the expression pattern in the liver, including metabolic key enzymes (e.g. *Dmdgh, Gnmt, Cth, Cyp2c37*)^24^. Several of these genes were implicated in mediating the benefits of caloric restriction^71–73^. For example, CTH (CGL) mediates the dietary restriction response in mice via H_2_S production^74^. Similarly, food intake restriction increases GNMT levels, promotes energy homeostasis, and increases healthy lifespan^73,75^. Altogether, our data demonstrate that Titan mice are responsive to diet intervention, and thus may represent a suitable translational mouse model to assess the impact of diet or other therapeutic interventions on manifestations of obesity^16,29,76^.

The poly-genetic Titan mouse model may present several advantages over other currently used mouse models of unhealthy obesity. Notably, no further genetic alterations or elimination of regulatory genes are needed to achieve the obesity phenotype in Titan mice. Most genetic rodent models of obesity are based on the disruption of the leptin signaling pathway, including the mouse Leptin-deficient (ob/ob), the mouse leptin receptor-deficient (db/db), the rat Zucker Diabetic Fatty and DahlS.Z-Lepr^fa^/Lepr^fa^ (DS/obese), the Koletsky rats carrying a nonsense mutation in the leptin receptor, or the POUND mice (C57BL/6NCrl-Lepr db-lb/Crl)^16,76,77^. A major disadvantage of these rodent models is that they do not closely reflect human obesity, which is multifactorial and not always affecting leptin signaling, thus rendering it difficult to translate findings in these rodents to human patients. Another advantage of the Titan mice, notably when studying the impact of obesity on lifespan phenotype, is that ERF diet intervention could be already assessed in 4-5 months Titan mice, whereas similar intervention would take much longer in other mouse models^29^

In conclusion, the Titan mouse line represents an important scientific achievement in breeding of mice, making Titan mice one of the biggest laboratory mouse lines ever described. The detailed genomic, epigenetic and phenotypic analyses described here strongly support the Titan line as a relevant model to study various aspects of interventions in unhealthy obesity. The present phenotypic analysis provides a strong basis for further characterization of the Titan model as a novel tool for studying and potentially developing pharmaceutical interventions targeting obesity and associated disorders.

## Materials and methods

### Animals and housing conditions

All procedures were performed in accordance with national and international guidelines and approved by our own institutional board (Animal Protection Board from the Leibniz Institute for Farm Animal Biology). At the German Mouse Clinic, mice were maintained in IVC cages with water and standard mouse chow according to the directive 2010/63/EU, German laws and GMC housing conditions (www.mouseclinic.de). All tests were approved by the responsible authority of the district government of Upper Bavaria.

At the FBN, the animals were maintained in a specific pathogen-free (SPF) environment with defined hygienic conditions at a temperature of 22.5±0.2°C, at least 40% humidity and a controlled light regime with a 12:12-h light-dark cycle. The mice were kept in Polysulfon-cages of 365 x 207 x 140 mm (H-Temp PSU, Type II L, Eurostandard, Tecniplast, Germany) and had free access to pellet concentrate and water. Mice were fed ad libitum using a standard breeding food (SBF, GE=16.7 MJ/kg, ME= 14.0 MJ/kg) including 12% fat, 27% protein and 61% carbohydrates (ssniff® M-Z autoclavable, Soest, V1124-300, Germany).

For the energy-reduced survival experiment, the mice were fed with a mouse maintenance energy-reduced diet (ERF, GE=15.7 MJ/kg, ME= 11.7 MJ/kg) characterized by a low energy density and high fiber contents including 10% fat, 21% protein and 69% carbohydrates (ssniff® M-H autoclavable, Soest, V1574-300, Germany). Mice were placed under the ERF diet starting at 12 weeks of age.

In both control and Titan lines, the litter size was standardized to 10 (5 male / 5 female pubs) immediately after birth and until weaning occurs at the age of 21 days. At the age of three weeks, males of different litters were grouped in a cage at three animals per cage. The first survival experiment was done with n=104 Titan males from generation 181 and n=96 control males originating from generation 192. The second survival experiment under the ERF diet started with n=103 Titan and n=102 control males, one generation later. Additionally, 27 Titan male full siblings pairs of generation 183 were divided into two contemporaneous groups and were fed with SBF or ERF after 12 weeks of age to generate the samples to analyze metabolic parameters. Individual weights of all animals were taken regularly every three weeks, starting with the age of 21 days. Hybrid mice were generated by crossing unselected control mice to Titan mice (Titan mother x control father or control mother x Titan father).

During the survival experiments, all included males were observed daily for their health condition. If physical impairments were detected, which would cause considerable suffering or substantial pain to the animals, they were sacrificed and such incidents were documented accordingly.

### Origins of the growth-selected strain and the control strain

We used mice of an unselected strain (FZTDU) as control and a strain selected for high body mass at day 42 of age (DU6/Titan), both bred at the Leibniz Institute of Farm Animal Biology (FBN), Dummerstorf, Germany.

The initial population of mice was created in 1969/1970 by crossing four outbred (NMRI orig., Han/NMRI, CFW, CF1) and four inbred (CBA/Bln, AB/Bln, C57BL/Bln, XVII/Bln) populations^7^. Mice of the control line FZTDU used in this experiment were mated randomly over about 192 generations with a population size of 100 to 200 mating pairs per generation, respectively. Four generations of the control line are generated yearly using a rotation procedure of Poiley (1960) to avoid inbreeding^78^.

The growth selection started in 1975, thus creating the Dummerstorf growth line DU6 (Titan) by selection for high body mass by sib selection in a conventional environment. In every generation, four generations per year, 60-80 mating pairs were made at the age of 63 ± 3 days^8,9^. The selection procedure as described above was maintained for 153 generations. Only seven males and seven females of the DU6 line belonging to generation 154 and 155 (year 2011-2012), respectively, were used as new founder animals after embryo transfer in a newly built animal facility. Of note, in 2011/2012, the Titan mouse lines (generations 154 and 155) were transferred into a new state-of-the-art pathogen-free animal facility and their diet composition changed at that point^12^. Over the entire term of the following five generations the new breeding population of at least 60 pairs of Titan DU6 mice was established^9^, taking care of an equal distribution of the alleles of the 14 founder animals. In opposition to the former sib selection in generation number 161, breeding value estimation started introducing a two-trait BLUP (**B**est Linear **U**nbiased **P**rediction) animal model for male and female body mass. The raising inbreeding coefficient in the selection line was then controlled by the ‘Optimal Genetic Contributions to the Next Generations’ method of Meuwissen^79^.

### Histology, plasma, DEXA – GMC

Importantly, because Titan mice are not inbred, we increased the number of animals from 15 (standard GMC animal number per group and sex for inbred mice) to 20, focusing on one sex only (males) to avoid sex-specific confounding factors.

Two cohorts of 20 DU6/Titan and 20 control (FZTDU) male mice were subjected to an extensive phenotypic screening at the German Mouse Clinic (GMC), including standardized phenotyping in e.g. the areas of energy metabolism, clinical chemistry, pathology^80^ (see also www.mouseclinic.de). The phenotypic tests were part of the GMC screen and were performed according to standardized protocols as described before^81–83^. Variations in protocols are specified. Depending on the test performed, animal numbers varied, as indicated in the respective Figure/Table.

Plasma clinical chemistry analyses included determination of blood lipid and glucose levels on freshly collected Li-heparin-plasma samples collected after overnight food withdrawal and measurement of a broad spectrum of parameters including ALAT and ALP activity, urea and bilirubin levels in samples collected in ad libitum fed state during final blood withdrawal. Analyses were performed using an AU480 clinical chemistry analyzer (Beckman-Coulter, Germany) as described previously. Samples collected during final blood withdrawal were frozen and stored at −80°C until the day of analysis.

The determination of the IL-6, TNFα, Insulin, leptin and FGF21 plasma levels was performed with a combined electrochemiluminescence multiplexed assay system (Meso Scale Discovery, MSD, Rockville, MD USA).

X-ray imaging was performed in an UltraFocus DXA system (Faxitron Bioptics, LLC) with automatic exposure control.

Macroscopic examination was performed in combination with histopathological analyses using hematoxylin and eosin (H&E) staining on formalin-fixed paraffin-embedded sections (3 µm) of tissues from 29 organs as described in www.mouseclinic.de/screens/pathology. Immunohistochemistry was carried out in a Leica Bond III (Leica Biosystems) automatic stainer. Heat-induced antigen retrieval was performed with citrate buffer (pH 6) for 30 minutes (AR9961; Bond TM Epitope Retrieval Solution; Leica Biosystems) in 2-μm-thick sections. For the identification of specific cells in the thymus, antibodies against CD3 (Clone SP7; ZYT-RBG024; Zytomed systems) and CD45R/B220 (Clone RA3-6B2; 550286; BD Pharmingen) were employed and the staining was detected with DAB chromogen. The slides were scanned using a Hamamatsu NanoZoomer 2.0HT digital scanner and analyzed by two independent pathologists using NDP.view2 software (Hamamatsu Photonics).

### Analysis of genetic differentiation

The genomes of the Dummerstorf mouse lines were sequenced in order to conduct genomic analyses which included the detection of line-specific patterns of genetic differentiation with respect to the control line FZTDU^9^. This was done by averaging the per-SNP genetic differentiation (Fst^84^) in 50Kb windows (sliding window mode with size=50Kb and step=25Kb) containing at least 10 SNPs using vcftools v0.1.13^85^. Per-window Fst scores were standardized into z-scores, in order to evaluate the data as deviations from the genomic mean (z-Fst). The X-chromosome was standardized separately. Regions of distinct genetic differentiation (RDD) were defined as highly differentiated windows in DU6 (top 15% of the z-Fst distribution) that also displayed low genetic differentiation (bottom 25% of the z-Fst distribution) in a) all remaining lines or b) the rest of the lines, except DU6P. Genes overlapping this windows were detected and used for enrichment analysis using WebGestaltR^86^ Genomic regions around genes of interest were visualized with Gviz ^87^.

### Histone acetylation analysis

Histone acetylation analysis was done as previously described^34,35,88^. Briefly, 25 mg of liver was used for acid extraction for histone. The resuspended samples were loaded on gels and were stained overnight followed by distaining with water. Gel pieces were processed for mass spectrometry analysis using heavy labeling of d6-deuterated acetic anhydride. Histone peptides were desalted using C18-StageTips, eluted in 80% acetonitrile/0.25% TFA, vacuum concentrated, reconstituted in 25 µL 0.1% TFA and stored at 4°C (short) or 20°C (long).

Desalted peptides (5µl) were injected and separated on an Ultimate 3000 RSLCnano (ThermoFisher) with a gradient from 3 to 32% acetonitrile in 0.1% formic acid over 40 min at 400 nL/min in a 25-cm analytical Aurora C18 nanocolumn (75μm ID 120 Å, 1.6 μm, Ion Opticks). The effluent from the HPLC was directly electrosprayed into a Q Exactive HF instrument operated in data-dependent mode to automatically switch between full-scan MS and MS/MS acquisition. Survey full-scan MS spectra (from m/z 250–1600) were acquired with resolution R=60,000 at m/z 400 (AGC target of 3×10^6^). The ten most intense peptide ions with charge states between 2 and 5 were sequentially isolated to a target value of 1×10^5^ and fragmented at 27% normalized collision energy. Typical mass spectrometric conditions were: spray voltage, 1.5 kV; no sheath and auxiliary gas flow; heated capillary temperature, 250°C; ion selection threshold, 33,000 counts.

Peak integration was performed with Skyline (Skyline-daily (64bit) 4.2.1.19004) using doubly o triply charged MS1 precursor ions. A list of all precursor ions can be found in the supplementary Table S1. Peaks were selected manually and the integrated peak values (Area) were exported as .csv file for further calculations. After peak integration, the data summarization and statistical analysis were performed in Excel and R^34^.

### Staining of triglycerides in liver tissues

Liver tissue samples embedded in Tissue Tek (Weckert, Kitzingen, Germany) were cryosectioned (8 μm thick) using a Leica CM3050 S (Leica, Bensheim, Germany) cryostat microtome. After fixation in 4% paraformaldehyde / PBS for 1 min at RT, the slides were stained in RedOil solution (1 mg/ml RedOil (#A12989, Alfa Aesar, Karlsruhe, Germany) in 60% Isopropanol) for 10 min and then washed three times in distilled water. The slides were further stained in hematoxylin for 5 minutes and washed for 3 min in fresh tap water before embedding with Aquatex (Roth, Germany) and dried overnight. The staining of the triglycerides was visualized with a Nikon Microphot-Fxa microscope (Nikon Instruments Europe B.V., The Netherlands) and an image analysis system (Nikon Digital Sight, DS-L2).

### RNAseq

For RNA extraction and library preparation, 50 mg of tissues (liver) were homogenized in Trizol (Thermo Fisher; cat. no. 15596026) and processed according to the manufacturer’s instructions. RNA concentration and A_260/280_ ratio were measured with NanoDrop, followed by Bioanalyzer using RNA pico assay kit using manufacturer’s protocol. rRNA depletion was performed using NEBNext rRNA Depletion Kit (Human/Mouse/Rat) [NEB #E6310] and library preparation for RNA-sequencing was performed using NEBNext Ultra II Directional RNA Library Prep Kit for Illumina [NEB #E7760] following manufacturer’s protocol. Libraries were sequenced on an Illumina HiSeq 1500 instrument at the Laboratory of Functional Genomic Analysis (LAFUGA, Gene Center Munich, LMU).

For RNA-seq data analysis, read mapping of mouse tissue samples to the mouse genome (GRCm38) and counting of reads mapped to genes were performed using STAR v2.5.3a^89^ using parameters --quantMode GeneCounts and providing annotation –sjdbGTFfile Mus_musculus.GRCm38.97.gtf. Aligned reads were filtered for unmapped, multi mapped and ambiguous reads. Reads from histones and Y chromosome were removed. Reads were also filtered if they had low read counts in at least two samples. Differential expression analysis was carried out using DESeq2 v1.24.0^90^ at an adjusted p-value cut-off of 0.05. GO term analysis was performed using ClusterProfiler v3.12.0^91^ at FDR of 0.05 using Benjamini-Hochberg procedure and with a log fold change cut-off of 0.5. GO terms containing at least a minimum of 10 genes were considered.

All the plots generated for RNA sequencing data were obtained using ggplot2 v3.2.1^92^ unless otherwise stated. For heatmaps and Venn diagram for RNA sequencing and proteomics data, pheatmap v1.0.12 (Kolde, R. (2013). pheatmap: Pretty Heatmaps. R package version 0.7.7. http://CRAN.R-project.org/package=pheatmap) were used respectively with genes (or proteins) passing the adjusted p-value significance of 0.05.

### Proteomics

The proteome protocol was adopted from^88,93^ with the following modifications. A total of 200 mg of frozen mice liver was homogenized in 500 µl lysis buffer [50 mM Tris-HCl pH 7.5, 500 mM NaCl, 1 ml EDTA, 0.1% NP-40 and 20% glycerol, 15 mM sodium butyrate, 60 mM of sirtinol and one protease inhibitor tablet (Roche)] and then supplemented with 200 µl 6 M urea/2 M thiourea and 900 µl lysis buffer. To reduce disulfide bonds, samples were treated with 1 mM DTT for 45 min at 4°C, followed by a treatment with 550 mM IAA for 30 min at 4°C in the dark. 1 M ammonium bicarbonate (Ambic) was added to the samples to get a final concentration of 1 M urea. The proteins were digested for 5 hours with Lys-C (Wako) at room temperature and overnight with trypsin (Worthington). Samples were acidified and diluted with TFA to a final concentration of 1% TFA before being loaded on the Sep-Pak Light C18 cartridges (Waters). Columns were washed with 0.1% TFA and eluted with 60% acetonitrile (ACN)/0.25% TFA. The eluates were dried out by speed vacuum. The pellets were re-dissolved in buffer [50 mM Hepes pH 8.0 and 50 mM NaCl] and the protein concentration was measured by Nanodrop.

LC-MS/MS measurements were performed on an Ultimate 3000 RSLCnano system coupled to a Q-Exactive HF-X mass spectrometer (ThermoFisher Scientific). For full proteome analyses, ∼0.25 µg of peptides were delivered to a trap column (ReproSil-pur C18-AQ, 5 μm, Dr. Maisch, 20 mm × 75 μm, self-packed) at a flow rate of 5 μL/min in 100% solvent A (0.1% formic acid in HPLC grade water). After 10 minutes of loading, peptides were transferred to an analytical column (ReproSil Gold C18-AQ, 3 μm, Dr. Maisch, 450 mm × 75 μm, self-packed) and separated using a 110 min gradient from 4% to 32% of solvent B (0.1% formic acid in acetonitrile and 5% (v/v) DMSO) at 300 nL/min flow rate. Both nanoLC solvents (solvent A = 0.1% formic acid in HPLC grade water and 5% (v/v) DMSO) contained 5% DMSO to boost MS intensity.

The Q-Exactive HF-X mass spectrometer was operated in data-dependent acquisition (DDA) and positive ionization mode. MS1 spectra (360–1300 m/z) were recorded at a resolution of 60,000 using an automatic gain control (AGC) target value of 3e6 and maximum injection time (maxIT) of 45 msec. Up to 18 peptide precursors were selected for fragmentation in the case of the full proteome analyses. Only precursors with charge state 2 to 6 were selected and dynamic exclusion of 30 sec was enabled. Peptide fragmentation was performed using higher energy collision-induced dissociation (HCD) and a normalized collision energy (NCE) of 26%. The precursor isolation window width was set to 1.3 m/z. MS2 resolution was 15,000 with an automatic gain control (AGC) target value of 1e^5^ and maximum injection time (maxIT) of 25 msec (full proteome).

Peptide identification and quantification were performed using MaxQuant (version 1.6.3.4) with its built-in search engine Andromeda^94,95^. MS2 spectra were searched against the Uniprot mus musculus proteome database (UP000000589, 54,208 protein entries, downloaded 22.3.2019) supplemented with common contaminants (built-in option in MaxQuant). Trypsin/P was specified as the proteolytic enzyme. Precursor tolerance was set to 4.5 ppm, and fragment ion tolerance to 20 ppm. Results were adjusted to 1% false discovery rate (FDR) on peptide spectrum match (PSM) level and protein level employing a target-decoy approach using reversed protein sequences. The minimal peptide length was defined as 7 amino acids, the “match-between-run” function was disabled. For full proteome analyses, carbamidomethylated cysteine was set as fixed modification and oxidation of methionine and N-terminal protein acetylation as variable modifications.

For the statistical analysis of the proteomics data, six biological replicates were measured in young and old Titan as well as young and old control mice. Protein abundances were calculated using the LFQ algorithm from MaxQuant^96^. Before further downstream analyses, protein LFQ values were logarithm (base 10) transformed. Next, Limma^97^ was used to identify the differentially expressed proteins between young control vs young Titan; young control vs old control; young Titan vs old Titan, and old control vs old Titan. The resulted p-values were adjusted by the Benjamini-Hochberg algorithm^98^ to control the false discovery rate (FDR). The differential analyses were performed on proteins that were identified in at least four out of six biological replicate samples in both groups under comparison.

Gene set annotations were downloaded from MSigDB^99^, including the Gene Ontology annotation (C5 category) and pathway annotation (C2 category). The gene IDs of differentially expressed proteins were mapped to the gene set annotations. The significance of over-representation was evaluated using fisher’s exact test.

### Total RNA extraction for quantitative RT-PCR

RNA extraction was performed from 50 mg deep-frozen liver tissue. The tissue was homogenized in 1 ml TRItidy G™ via pestle and incubated at room temperature for 5 minutes. After the addition of 200 µl chloroform, the sample was mixed 15 seconds with the Vortex and again incubated for 2 min at room temperature with a following centrifugation step at 12,000 x g, 4°C for 15 min. The aqueous phase was transferred into a new vial and mixed with 500 µl - 20°C isopropanol and incubated 10 minutes at −20°C. The samples were then centrifuged at 8,000 x g, 4°C for 10 minutes and the supernatant was discarded. Two washing steps with 1 ml −20°C cold 70% ethanol were applied and the pellet was air-dried and dissolved in 100 µl nuclease-free water. RNA concentration was measured with NanoDrop 2000.

### Quantitative Real-time PCR

A total of 800 ng RNA was used for cDNA synthesis, using the SensiFAST™cDNA Synthesis Kit (Bioline). Quantitative PCR was performed with the SensiFAST™SYBR® No-Rox Kit (Bioline) using 1.25 ng per reaction, except for the testing of mTOR (2.5 ng), and 4 pmol of forward and reverse primers. Quantitative PCR (qPCR) reactions were run on the Roche Lightcycler 96. The annealing temperature was 60°C. The qPCR data were normalized to the levels of beta-actin. Primers are listed in Table S5. PCR products were cleaned up using the High Pure PCR Product Purification Kit (Roche) and sequenced by LGC Genomics GmbH. Each sample was measured using two technical replicates.

### Statistics and visualization

Unless stated otherwise in the RNA-seq and proteomics experimental method, statistics and graphing were conducted on GraphPad Prism 9. Unpaired two-tailed *t*-tests with Welch’s correction (parametric) or MWU testing (non-parametric) were used for calculating the *P*-values unless stated otherwise. A multiple correction (FDR) was performed for SBF vs ERF qPCR and plasma analysis. A p-value ≤0.05 was used as level of significance.

Importantly, according to the GMC analysis outline, correction for multiple testing was not performed when analyzing the GMC data set (See GMC website).

## Data availability

The proteomic raw data and MaxQuant search files have been deposited to the ProteomeXchange Consortium (http://proteomecentral.proteomexchange.org) via the PRIDE partner repository and can be accessed using the dataset identifier PXD019030 (reviewer account details: username; password rvqDlf58). All RNAseq data have been deposited to GEO.

## Acknowledgments

We would like to thank our technician Verena Hofer-Pretz for performing many of the experiments for this study, as well as managing the laboratory conditions that enabled this work. The authors thank Ines Müntzel, Karin Ullerich, Sabine Maibohm, Benita Lucht, Hildburg Meier and Diane Loth from the mouse facility for the animal care and technical assistance. The authors also thank Erika Wytrwat for technical support. The GMC would like to thank the technicians and animal care takers involved in this project for their expert technical help. We also thank Franziska Hackbarth, Nina Lomp and Hermine Kienberger for their excellent laboratory assistance at the BayBioMS. The authors thank Dr. Anne Rascle of AR Medical Writing (Regensburg, Germany) for providing medical writing and editing support. The authors also thank Dario Riccardo Valenzano and Christoph Anacker for their comments on the manuscript.

The study was supported by the Leibnitz society and by the German Federal Ministry of Education and Research (Infrafrontier grant 01KX1012 to MHdA); German Center for Diabetes Research (DZD) (MHdA), the Helmholtz Alliance ‘Aging and Metabolic Programming, AMPr’ (RG).

## Author contributions

SP conceptualized the project. ML prepared all the mice work, conducted the survival curves with the weight measurements and assisted with conceptualizing this work. ML also maintains the Dummerstorf selection line. AH contributed to this work by the management of pilot studies on life expectancy in the mouse model used in this study. ID-D extracted the RNA for RNAseq. AV-V prepared the libraries for transcriptomic analysis. AM-E prepared the proteomics and histone acetylation. AS-M, RG, BR, AA-P, TK-R, JC-W performed mouse phenotyping tests and analyzed the data, LB supported data analyzes and manuscript preparation, CS coordinated and VG-D, HF and MHdA supervised the project at the GMC and conceived phenotyping test pipeline. AV-V analyzed the RNA-seq and histone acetylation and generated the data visualization. AI supervised AV-V’s work. SPV and JS conducted and analyzed the RDD data. CM and CL ran, analyzed and visualized the proteome data. FK isolated the blood for the ERF experiments. VC ad JB prepared and analyzed the Oil red O/ hematoxylin straining. ID-D, FC, AK and SP wrote the initial manuscript. All authors contributed to the final version of the manuscript. AM-E generated all the final figures for the paper. BG assisted with graph analysis and manuscript preparation.

## Supplementary figures

**Figure S1. Titan mice display several molecular criteria for MUO.**

(A-C) Plasma analyses comparing control and Titan mice after fasting at 18–20 weeks. Titan mice showed higher levels of triglycerides (p = 0.0055) and both non-HDL (p = 0.0022) and HDL cholesterol (n= 20 control vs 19 Titan). (D) Insulin levels in control (n = 19) and Titan (n = 18) mice at 16–17 weeks of age. (E) Leptin levels in control (n = 20) and Titan (n = 18) mice and (F) higher FGF21 levels in the Titan mice line compared to the control mice line (N=20 control vs 18 Titan). \*\**P* < 0.01 \*\*\**P* < 0.001. Error bars indicate SEMs. Unpaired two-tailed *t-*tests with Welch’s correction were used to calculate *P-*values.

**Figure S2. Titan mice show signs of heart fibrosis.**

H&E and Sirius Red stainings of the heart from 16-17-weeks-old control and Titan mice are shown (0.45x magnification). Sirius Red staining allows a better visualization of fibrotic tissue. Higher magnification pictures from rectangles were taken at 10x.

**Figure S3. Titan mice show thickened dermis and early signs of increased inflammation.**

Representative images of hematoxylin and eosin (H&E) staining of the skin. Comparison of Titan and control mice at 16–17 weeks of age. (n = 4–6 per group). (A) The dermis of the Titan mice is substantially thicker compared to control mice. (B and C) IL-6 plasma levels (p = 0.0086) in control (n = 13) and Titan (n = 17) mice, and TNFα plasma levels in control (n = 20) and Titan (n = 18) mice. (D) Representative images of hematoxylin and eosin (H&E) staining and B-cell immunohistochemistry (IHC) of the thymus of control (left) and Titan (right) mice (2.5x magnification). IHC of thymic medullar nodes revealed that they are composed mainly of B-cells (CD45R/B220-positive) instead of T-cells (CD3-positive). Images in squares were taken at 10x magnification (n = 4–6 per group). \**P* < 0.05, \*\*\**P* < 0.001. Error bars indicate SEMs. Unpaired two-tailed *t-*tests with Welch’s correction were used to calculate *P-*values.

**Figure S4. Liver H&E staining comparison of control vs Titan mice.**

Shown are representative H&E-stained images of the liver from 16-17-weeks old control and Titan mice (10x and 40x magnifications are shown).

**Figure S5. Transcriptome changes in the Liver of younger vs older Titan mice.**

(A) Heat map showing significantly altered genes between 11-week-old control and Titan mice. (B) Heat map showing significantly altered genes between 11- and 19-21-week-old Titan mice. (C) GO term analysis of (B) revealed an increase in insulin-related pathway and glucose homeostasis and a decrease in various metabolic processes including fatty acid metabolism. (D) Heat map showing significantly altered genes between 19-21-week-old control and Titan mice. n=5 per group.

**Figure S6. Liver proteome analyses reveal differences in metabolic protein levels between control and Titan mice.**

(A) Heat map comparing proteomes of 11-week-old control and Titan mice.

(B) Heat map comparing proteomes of 19-21-week-old control and Titan mice.

(C) GO term analysis of 11-week-old mice indicated increased fatty acid metabolism, various lipid catabolism, and coenzyme metabolism while downregulation in xenobiotic, amino acid and sulfur metabolism. (D) GO term analysis of 19-21-week-old mice showed upregulation of proteins involved in fatty acid metabolism, carbohydrate catabolism and coenzyme metabolism and downregulation of proteins involved in amino acid and sulfur metabolism (n = 6 per group).

**Table S1. Unique RDD in Titan mice**

**Table S2. Unique RDD in Titan mice excluding DU6P mice**

**Table S3.**
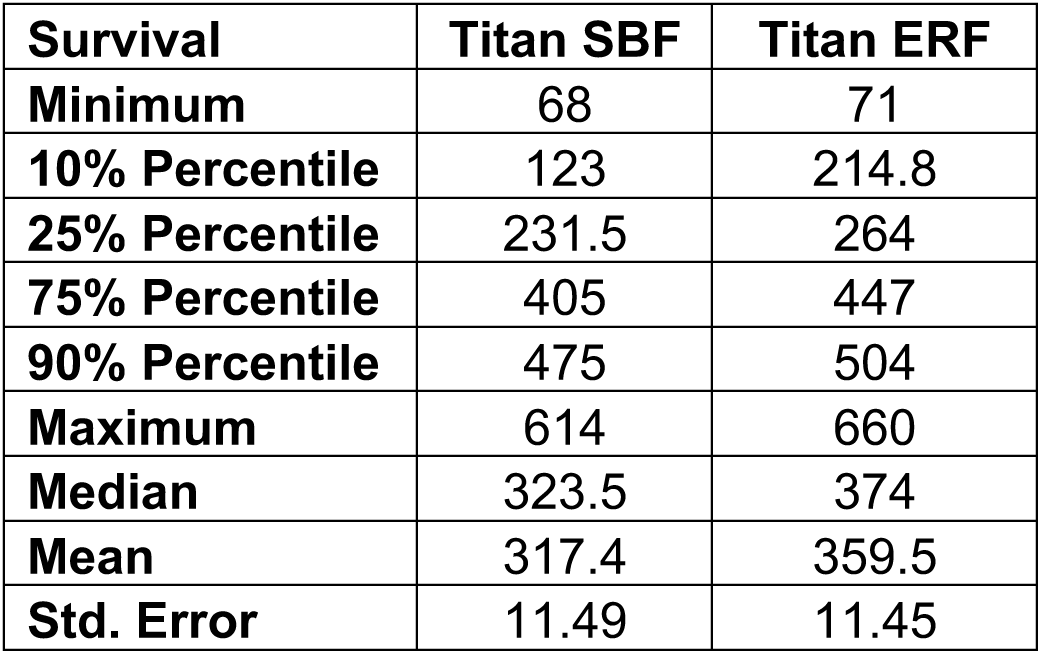
Lifespans in days for Titan mice receiving SBF and ERF, respectively.

| **Survival** | **Titan SBF** | **Titan ERF** |
| --- | --- | --- |
| **Minimum** | 68 | 71 |
| **10% Percentile** | 123 | 214.8 |
| **25% Percentile** | 231.5 | 264 |
| **75% Percentile** | 405 | 447 |
| **90% Percentile** | 475 | 504 |
| **Maximum** | 614 | 660 |
| **Median** | 323.5 | 374 |
| **Mean** | 317.4 | 359.5 |
| **Std. Error** | 11.49 | 11.45 |

**Tables S4-5. List of significantly altered genes (RNAseq)**

**Tables S6. Proteome**

**Table S7. Key resources and Real-time PCR primer sequences.**

## References

1. Bolker, J. A. Animal Models in Translational Research: Rosetta Stone or Stumbling Block? Bioessays 39, 1700089 (2017).

2. Choudhary, A. & Ibdah, J. A. Animal models in today’s translational medicine world. Mo Med 110, 220–2 (2013).

3. Prabhakar, S. Translational Research Challenges. J Invest Med 60, 1141 (2012).

4. Brockmann, G. A. & Bevova, M. R. Using mouse models to dissect the genetics of obesity. Trends Genet 18, 367–376 (2002).

5. Renne, U. et al. Lifelong obesity in a polygenic mouse model prevents age- and diet-induced glucose intolerance-obesity is no road to late-onset diabetes in mice. PloS one 8, e79788 (2013).

6. Dietl, G., Langhammer, M. & Renne, U. Model simulations for genetic random drift in the outbred strain Fzt:DU. Arch Anim Breed 47, 595–604 (2004).

7. Schüler, L. Selection for fertility in mice - the selection plateau and how to overcome it. TAG. Theoretical and applied genetics. Theoretische und angewandte Genetik 70, 72–79 (1985).

8. Bünger, L., Renne, U., Dietl, G. & Kuhla, S. Long-term selection for protein amount over 70 generations in mice. Genetical research 72, 93–109 (1998).

9. Palma-Vera, S. E. et al. Genomic characterization of world’s longest selection experiment in mouse reveals the complexity of polygenic traits. (2021) doi:10.1101/2021.05.28.446207.

10. Timtchenko, D. et al. Fat storage capacity in growth-selected and control mouse lines is associated with line-specific gene expression and plasma hormone levels. International journal of obesity and related metabolic disorders : journal of the International Association for the Study of Obesity 23, 586–594 (1999).

11. Aksu, S., Koczan, D., Renne, U., Thiesen, H.-J. & Brockmann, G. A. Differentially expressed genes in adipose tissues of high body weight-selected (obese) and unselected (lean) mouse lines. Journal of applied genetics 48, 133–143 (2007).

12. Brenmoehl, J. et al. Partial phenotype conversion and differential trait response to conditions of husbandry in mice. J Comp Physiology B 188, 527–539 (2018).

13. Walz, M. et al. Overlap of Peak Growth Activity and Peak IGF-1 to IGFBP Ratio: Delayed Increase of IGFBPs Versus IGF-1 in Serum as a Mechanism to Speed up and down Postnatal Weight Gain in Mice. Cells 9, 1516 (2020).

14. Iacobini, C., Pugliese, G., Fantauzzi, C. B., Federici, M. & Menini, S. Metabolically healthy versus metabolically unhealthy obesity. Metabolis 92, 51–60 (2019).

15. Shinohara, M. et al. Increased levels of Aβ42 decrease the lifespan of ob/ob mice with dysregulation of microglia and astrocytes. Faseb J 34, 2425–2435 (2020).

16. Kennedy, A. J., Ellacott, K. L. J., King, V. L. & Hasty, A. H. Mouse models of the metabolic syndrome. Disease models & mechanisms 3, 156–166 (2010).

17. Guilherme, A., Henriques, F., Bedard, A. H. & Czech, M. P. Molecular pathways linking adipose innervation to insulin action in obesity and diabetes mellitus. Nature reviews. Endocrinology 15, 207–225 (2019).

18. Lund, J., Lund, C., Morville, T. & Clemmensen, C. The unidentified hormonal defense against weight gain. PLoS biology 18, e3000629 (2020).

19. Giralt, M., Gavaldà-Navarro, A. & Villarroya, F. Fibroblast growth factor-21, energy balance and obesity. Mol Cell Endocrinol 418, 66–73 (2015).

20. Zhang, X. et al. Serum FGF21 levels are increased in obesity and are independently associated with the metabolic syndrome in humans. Diabetes 57, 1246–1253 (2008).

21. Cavalera, M., Wang, J. & Frangogiannis, N. G. Obesity, metabolic dysfunction, and cardiac fibrosis: pathophysiological pathways, molecular mechanisms, and therapeutic opportunities. Transl Res 164, 323–335 (2014).

22. Shimizu, I. & Walsh, K. The Whitening of Brown Fat and Its Implications for Weight Management in Obesity. Curr Obes Reports 4, 224–229 (2015).

23. Selman, C., Nussey, D. H. & Monaghan, P. Ageing: it’s a dog’s life. Current biology : CB 23, R451–3 (2013).

24. Toye, A. A. et al. Subtle metabolic and liver gene transcriptional changes underlie diet-induced fatty liver susceptibility in insulin-resistant mice. Diabetologia 50, 1867–1879 (2007).

25. Fontana, L. & Partridge, L. Promoting health and longevity through diet: from model organisms to humans. Cell 161, 106–118 (2015).

26. Francesco, A. D., Germanio, C. D., Bernier, M. & Cabo, R. de. A time to fast. Science (New York, N.Y.) 362, 770–775 (2018).

27. Greer, E. L. & Brunet, A. Different dietary restriction regimens extend lifespan by both independent and overlapping genetic pathways in C. elegans. Aging cell 8, 113–127 (2009).

28. Bastías-Pérez, M., Serra, D. & Herrero, L. Dietary Options for Rodents in the Study of Obesity. Nutrients 12, 3234 (2020).

29. Li, H. & Auwerx, J. Mouse Systems Genetics as a Prelude to Precision Medicine. Trends in genetics : TIG (2020) doi:10.1016/j.tig.2020.01.004.

30. Liao, C.-Y., Rikke, B. A., Johnson, T. E., Diaz, V. & Nelson, J. F. Genetic variation in the murine lifespan response to dietary restriction: from life extension to life shortening. Aging cell 9, 92–95 (2010).

31. Derraik, J. G. B. et al. Effects of Age, Gender, BMI, and Anatomical Site on Skin Thickness in Children and Adults with Diabetes. Plos One 9, e86637 (2014).

32. Obri, A., Serra, D., Herrero, L. & Mera, P. The role of epigenetics in the development of obesity. Biochem Pharmacol 177, 113973 (2020).

33. Mutlu, B. & Puigserver, P. GCN5 ACETYLTRANSFERASE IN CELLULAR ENERGETIC AND METABOLIC PROCESSES. Biochimica Et Biophysica Acta Bba - Gene Regul Mech 1864, 194626 (2020).

34. Feller, C., Forne, I., Imhof, A. & Becker, P. B. Global and specific responses of the histone acetylome to systematic perturbation. Molecular cell 57, 559–571 (2015).

35. Bux, E. M. et al. Determining histone H4 acetylation patterns in human peripheral blood mononuclear cells using mass spectrometry. Clin Mass Spectrom 15, 54–60 (2019).

36. Lerin, C. et al. GCN5 acetyltransferase complex controls glucose metabolism through transcriptional repression of PGC-1α. Cell Metab 3, 429–438 (2006).

37. Struhl, K. Histone acetylation and transcriptional regulatory mechanisms. Gene Dev 12, 599–606 (1998).

38. Giannini, E. G., Testa, R. & Savarino, V. Liver enzyme alteration: a guide for clinicians. Can Med Assoc J 172, 367–379 (2005).

39. Mews, P. et al. Acetyl-CoA synthetase regulates histone acetylation and hippocampal memory. Nature (2017) doi:10.1038/nature22405.

40. Yu, H., Pardoll, D. & Jove, R. STATs in cancer inflammation and immunity: a leading role for STAT3. Nat Rev Cancer 9, 798–809 (2009).

41. Heinrich, P. C. et al. Principles of interleukin (IL)-6-type cytokine signalling and its regulation. The Biochemical journal 374, 1–20 (2003).

42. Landskron, G., Fuente, M. D. la, Thuwajit, P., Thuwajit, C. & Hermoso, M. A. Chronic inflammation and cytokines in the tumor microenvironment. Journal of immunology research 2014, 149185 (2014).

43. Bao, P., Liu, G. & Wei, Y. Association between IL-6 and related risk factors of metabolic syndrome and cardiovascular disease in young rats. International journal of clinical and experimental medicine 8, 13491–13499 (2015).

44. Popko, K. et al. Proinflammatory cytokines Il-6 and TNF-α and the development of inflammation in obese subjects. European journal of medical research 15 Suppl 2, 120–122 (2010).

45. Lee, J. H. et al. An engineered FGF21 variant, LY2405319, can prevent non-alcoholic steatohepatitis by enhancing hepatic mitochondrial function. Am J Transl Res 8, 4750–4763 (2016).

46. Morinaga, H. et al. Characterization of Distinct Subpopulations of Hepatic Macrophages in HFD/Obese Mice. Diabetes 64, 1120–1130 (2014).

47. Weisberg, S. P. et al. Obesity is associated with macrophage accumulation in adipose tissue. J Clin Invest 112, 1796–1808 (2003).

48. Nuñez, A., Rodrigo-Angulo, M. L., Andrés, I. D. & Garzón, M. Hypocretin/Orexin Neuropeptides: Participation in the Control of Sleep-Wakefulness Cycle and Energy Homeostasis. Curr Neuropharmacol 7, 50–59 (2009).

49. Tan, Y. et al. Impaired hypocretin/orexin system alters responses to salient stimuli in obese male mice. J Clin Invest 130, 4985–4998 (2020).

50. Valenzano, D. R. et al. The African Turquoise Killifish Genome Provides Insights into Evolution and Genetic Architecture of Lifespan. Cell 163, 1539–1554 (2015).

51. Díaz-Delfín, J. et al. TNF-α Represses β-Klotho Expression and Impairs FGF21 Action in Adipose Cells: Involvement of JNK1 in the FGF21 Pathway. Endocrinology 153, 4238–4245 (2012).

52. Grundy, S. M. et al. Diagnosis and management of the metabolic syndrome: an American Heart Association/National Heart, Lung, and Blood Institute scientific statement: Executive Summary. Critical pathways in cardiology 4, 198–203 (2005).

53. Pezzilli, R. & Calculli, L. Pancreatic steatosis: Is it related to either obesity or diabetes mellitus? World J Diabetes 5, 415 (2014).

54. Pinnick, K. E. et al. Pancreatic Ectopic Fat Is Characterized by Adipocyte Infiltration and Altered Lipid Composition. Obesity 16, 522–530 (2008).

55. Vidra, N., Trias-Llimós, S. & Janssen, F. Impact of obesity on life expectancy among different European countries: secondary analysis of population-level data over the 1975-2012 period. Bmj Open 9, e028086 (2019).

56. Fontaine, K. R., Redden, D. T., Wang, C., Westfall, A. O. & Allison, D. B. Years of Life Lost Due to Obesity. Jama 289, 187–193 (2003).

57. Jura, M. & Kozak, Leslie. P. Obesity and related consequences to ageing. Age 38, 23 (2016).

58. Frasca, D. & Blomberg, B. B. Adipose tissue, immune aging, and cellular senescence. Semin Immunopathol 42, 573–587 (2020).

59. Monteiro, R. & Azevedo, I. Chronic inflammation in obesity and the metabolic syndrome. Mediators of inflammation 2010, (2010).

60. Sun, S., Ji, Y., Kersten, S. & Qi, L. Mechanisms of Inflammatory Responses in Obese Adipose Tissue. Annu Rev Nutr 32, 261–286 (2012).

61. Langhammer, M. et al. Two mouse lines selected for large litter size display different lifetime fecundities. Reproduction (2021) doi:10.1530/rep-20-0563.

62. Lewis, D. F. V. 57 varieties: the human cytochromes P450. Pharmacogenomics 5, 305–318 (2004).

63. Morgan, E. T. Impact of infectious and inflammatory disease on cytochrome P450-mediated drug metabolism and pharmacokinetics. Clinical pharmacology and therapeutics 85, 434–438 (2009).

64. Siewert, E. et al. Hepatic cytochrome P450 down-regulation during aseptic inflammation in the mouse is interleukin 6 dependent. Hepatology (Baltimore, Md.) 32, 49–55 (2000).

65. Crocco, P. et al. Inter-Individual Variability in Xenobiotic-Metabolizing Enzymes: Implications for Human Aging and Longevity. Genes 10, (2019).

66. Peleg, S., Feller, C., Ladurner, A. G. & Imhof, A. The Metabolic Impact on Histone Acetylation and Transcription in Ageing. Trends in biochemical sciences 41, 700–711 (2016).

67. Bradshaw, P. C. Acetyl-CoA Metabolism and Histone Acetylation in the Regulation of Aging and Lifespan. Antioxidants 10, 572 (2021).

68. Karpe, F., Dickmann, J. R. & Frayn, K. N. Fatty Acids, Obesity, and Insulin Resistance: Time for a Reevaluation. Diabetes 60, 2441–2449 (2011).

69. Mahendran, Y. et al. Glycerol and Fatty Acids in Serum Predict the Development of Hyperglycemia and Type 2 Diabetes in Finnish Men. Diabetes Care 36, 3732–3738 (2013).

70. Wilkinson, M. J. et al. Ten-Hour Time-Restricted Eating Reduces Weight, Blood Pressure, and Atherogenic Lipids in Patients with Metabolic Syndrome. Cell Metab 31, 92–104.e5 (2019).

71. Chou, M. W., Kong, J., Chung, K. T. & Hart, R. W. Effect of caloric restriction on the metabolic activation of xenobiotics. Mutation research 295, 223–235 (1993).

72. Hine, C. et al. Endogenous hydrogen sulfide production is essential for dietary restriction benefits. Cell 160, 132–144 (2015).

73. Obata, F. & Miura, M. Enhancing S-adenosyl-methionine catabolism extends Drosophila lifespan. Nature communications 6, 8332 (2015).

74. Bithi, N. et al. Dietary restriction transforms the mammalian protein persulfidome in a tissue-specific and cystathionine γ-lyase-dependent manner. Nat Commun 12, 1745 (2021).

75. Obata, F. et al. Necrosis-driven systemic immune response alters SAM metabolism through the FOXO-GNMT axis. Cell reports 7, 821–833 (2014).

76. Hattori, T. et al. Characterization of a new animal model of metabolic syndrome: the DahlS.Z-Leprfa/Leprfa rat. Nutr Diabetes 1, e1–e1 (2011).

77. Wong, S. K., Chin, K.-Y., Suhaimi, F. H., Fairus, A. & Ima-Nirwana, S. Animal models of metabolic syndrome: a review. Nutr Metabolism 13, 65 (2016).

78. Poiley, S. M. A systematic method of breeder rotation for non-inbred laboratory animal colonies. Proc Anim Care Panel 159–166 (1960).

79. Meuwissen, T. H. Maximizing the response of selection with a predefined rate of inbreeding. Journal of animal science 75, 934–940 (1997).

80. Gailus-Durner, V. et al. Introducing the German Mouse Clinic: open access platform for standardized phenotyping. Nature methods 2, 403–404 (2005).

81. Fuchs, H. et al. Mouse phenotyping. Methods (San Diego, Calif.) 53, 120–135 (2011).

82. Rathkolb, B. et al. Clinical Chemistry and Other Laboratory Tests on Mouse Plasma or Serum. Current protocols in mouse biology 3, 69–100 (2013).

83. Rozman, J. et al. Glucose tolerance tests for systematic screening of glucose homeostasis in mice. Current protocols in mouse biology 5, 65–84 (2015).

84. Weir, B. S. & Cockerham, C. C. ESTIMATING F-STATISTICS FOR THE ANALYSIS OF POPULATION STRUCTURE. Evolution 38, 1358–1370 (1984).

85. Danecek, P. et al. The variant call format and VCFtools. Bioinformatics 27, 2156–2158 (2011).

86. Liao, Y., Wang, J., Jaehnig, E. J., Shi, Z. & Zhang, B. WebGestalt 2019: gene set analysis toolkit with revamped UIs and APIs. Nucleic Acids Res 47, W199–W205 (2019).

87. Hahne, F. & Ivanek, R. Statistical Genomics, Methods and Protocols. Methods Mol Biology 1418, 335–351 (2016).

88. Peleg, S. et al. Life span extension by targeting a link between metabolism and histone acetylation in Drosophila. EMBO reports 17, 455–469 (2016).

89. Dobin, A. & Gingeras, T. R. Mapping RNA-seq Reads with STAR. Current protocols in bioinformatics 51, 11.14.1–11.14.19 (2015).

90. Love, M. I., Huber, W. & Anders, S. Moderated estimation of fold change and dispersion for RNA-seq data with DESeq2. Genome biology 15, 550 (2014).

91. Yu, G., Wang, L.-G., Han, Y. & He, Q.-Y. clusterProfiler: an R package for comparing biological themes among gene clusters. Omics : a journal of integrative biology 16, 284–287 (2012).

92. Wickham, H. ggplot2, Elegant Graphics for Data Analysis. 147–168 (2016) doi:10.1007/978-3-319-24277-4_7.

93. Gaucher, J. et al. Distinct metabolic adaptation of liver circadian pathways to acute and chronic patterns of alcohol intake. Proceedings of the National Academy of Sciences of the United States of America 116, 25250–25259 (2019).

94. Cox, J. et al. Andromeda: a peptide search engine integrated into the MaxQuant environment. Journal of proteome research 10, 1794–1805 (2011).

95. Tyanova, S., Temu, T. & Cox, J. The MaxQuant computational platform for mass spectrometry-based shotgun proteomics. Nature protocols 11, 2301–2319 (2016).

96. Cox, J. et al. Accurate proteome-wide label-free quantification by delayed normalization and maximal peptide ratio extraction, termed MaxLFQ. Molecular & cellular proteomics : MCP 13, 2513–2526 (2014).

97. Ritchie, M. E. et al. limma powers differential expression analyses for RNA-sequencing and microarray studies. Nucleic acids research 43, e47 (2015).

98. Benjamini, Y. & Hochberg, Y. Controlling the False Discovery Rate: A Practical and Powerful Approach to Multiple Testing. Journal of the Royal Statistical Society 57, 289–300 (1995).

99. Liberzon, A. et al. The Molecular Signatures Database (MSigDB) hallmark gene set collection. Cell systems 1, 417–425 (2015).

